# An ERK-dependent molecular switch antagonizes fibrosis and promotes regeneration in spiny mice (*Acomys*)

**DOI:** 10.1101/2022.06.27.497314

**Authors:** Antonio Tomasso, Kerstin Bartscherer, Ashley W. Seifert

## Abstract

Although most mammals heal wounds with scar tissue, spiny mice (*Acomys cahirinus*) naturally regenerate skin and complex musculoskeletal tissues. The core signaling pathways driving mammalian tissue regeneration are poorly characterized. Here, we show that MAPK/ERK signaling acts as a major hub directing cellular injury responses towards regeneration. While immediate ERK activation was a shared feature of scarring (*Mus musculus*) and regenerating (*Acomys*) wounds, ERK activity was only sustained during regeneration. Upon ERK inhibition, the *Acomys* wound microenvironment exhibited key hallmarks of fibrosis, including a pro-scarring matrisome, epidermal differentiation and reduced proliferation. These findings indicate that ERK inhibition shifts tissue regeneration towards fibrosis. Conversely, the ectopic release of ERK activators (FGF/Neuregulin) in fibrotic scars, stimulated a pro-regenerative matrisome, cell proliferation and hair follicle neogenesis, thus promoting skin regeneration. Our data provide new insights into why some mammals regenerate better than others and open avenues to reverse fibrosis in favor of regeneration.

**Teaser:** FGF2 and NRG1 stimulate skin and hair follicle regeneration through ERK-mediated control of cell behavior in adult mammals.

## Introduction

Tissue damage or loss poses an immediate risk to organismal homeostasis. While the generation of scar tissue is a common response to repair wounded tissue in arthropods and vertebrates, some animals can completely regenerate lost tissue. A paradox is the seemingly haphazard distribution of this regenerative response across major metazoan clades (*1*). Moreover, even in closely related species, tissue damage can elicit different healing outcomes. For instance, the ability to regenerate a tail varies widely among reptiles, while the mechanistic basis for this variation remains unknown (*2*). Among mammals, spiny mice (*Acomys* species) naturally regenerate complex tissues in the ear pinna which includes full-thickness skin, hair follicles, adipose tissue, peripheral nerves, skeletal muscle and cartilage. In contrast, laboratory mouse strains (*Mus* species) and other murid rodents heal identical wounds with scar tissue characterized by excessive collagen deposition (*3-5*). Despite progress disentangling the mechanistic basis for these different healing responses, important questions remain. Tantamount among them is how the cellular response to injury is differentially regulated in regenerative and fibrotic healing.

Vertebrate regeneration occurs as a complex series of overlapping processes where inflammation and wound closure are followed by inflammatory resolution and the formation of a blastema, namely a mass of proliferating progenitor cells (*6*). Blastemal cells ultimately undergo morphogenesis to replace the damaged or missing tissue. The degree to which healing initiators, activated in response to injury, directly influence blastema formation remains unknown. Because enhanced regenerative ability has likely evolved *de novo* in mammals (*7*), comparing the healing response of identical injuries in regenerating and non-regenerating mammals offers an opportunity to identify cell autonomous drivers of mammalian regeneration. Given the highly conserved role that ERK-mediated signaling plays during the metazoan injury response (*8-10*), it is an attractive target to explore how it may control cell behavior during different modes of healing. Recent work has demonstrated that MAPK/ERK-mediated wound signals jumpstart the regenerative response in planarian flatworms (*11*).

Tissue regeneration occurs at a relatively slow rate in mammals and salamanders compared to invertebrates and fish, providing an opportunity to dissect how temporal ERK activation regulates the cellular response to injury throughout wound healing and blastema formation. By analyzing ERK activation during fibrotic repair in *Mus* and regenerative healing in *Acomys*, we show that (1) immediate ERK activation is a conserved feature of the injury response in regenerators (*Acomys*) and poor-regenerators (*Mus*), (2) ERK activation is sustained after wound closure only in *Acomys* during blastema formation where it is required to activate a cellular regeneration program in a tissue-specific manner, and (3) ectopic ERK activation confers regenerative traits to normally non-regenerating *Mus* tissue. Together, our data demonstrate that ERK activity acts as a molecular switch between regeneration and scarring and explains why even closely related mammalian species may exert very different cellular responses to injury.

## Results

### Injury-induced ERK activation is a common feature of wound healing in non-regenerating *Mus* and regenerating *Acomys*

To test our hypothesis that ERK activity might differentially regulate healing outcomes in *Acomys* and *Mus*, we first explored wound-induced ERK activation (phosphorylated ERK - pERK) using a 4-mm ear punch injury (*4, 5*) and a pERK-specific antibody (Fig. 1A and fig. S1, A and B). In relation to the long axis of the ear pinna, the ear punch generated a proximal and a distal segment where healing occurs (Fig. 1A). While active ERK was nearly absent in uninjured tissue (the exception being a few hair follicle bulb cells in both species), as early as 10 minutes after injury, we observed high levels of active ERK in both *Mus* and *Acomys* (Fig. 1, B to D). Multiple cell types, including keratinocytes, chondrocytes, perichondral cells, endothelial cells and mesenchymal cells immediately activated ERK at the wound site (Fig. 1C). Compared to *Mus*, ERK was more broadly activated in *Acomys* from 1h post-injury (hpi) (Fig. 1, E to G) and peaked at 6hpi in different cell populations (Fig. 1G and fig. S1, C and D). We also observed a bias in pERK^+^ cell types which delineated a spatio-temporal wave of ERK activation: the percentage of pERK^+^ nuclei was higher among keratinocytes in the epidermis and perichondral cells compared to chondrocytes in the cartilage (Fig. 1G and fig. S1, C and D). Hence, multiple cell lineages detected injury signals from the wound margin in the proximal and distal ear segment during the early phase of the injury response (fig. S1, E to H). Together, these data show that pERK acts as an immediate sensor of tissue damage in regeneration-incompetent (*Mus*) and -competent mammals (*Acomys*).

**Fig. 1.**
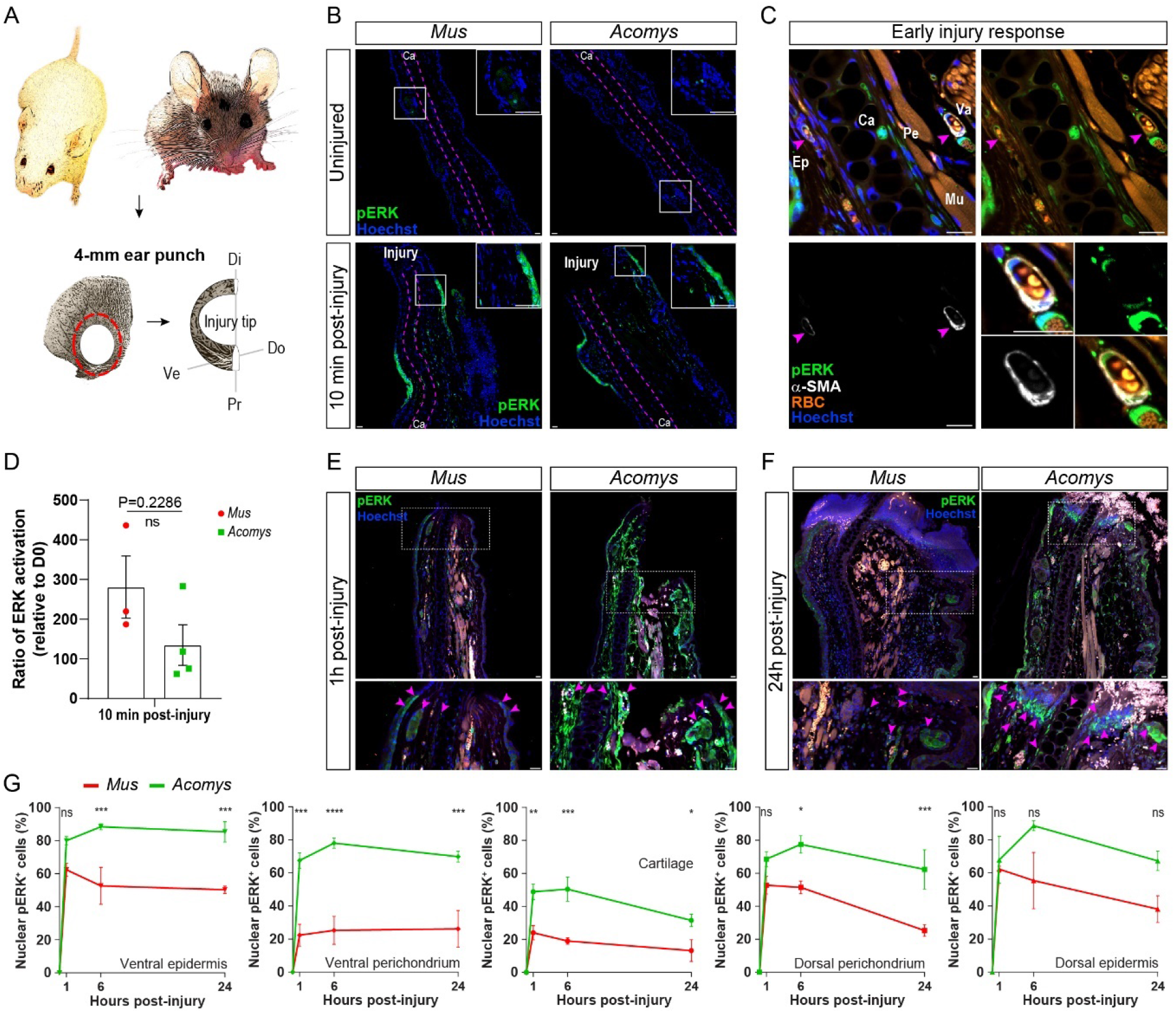
ERK activation is an immediate cellular response to wounds in fibrotic and regenerative healing. **(A)** Comparative studies between *Mus* (left) and *Acomys* (right). Schematics of the ear structures after ear punch: proximal segment (Pr); distal segment (Di), ventral (Ve) and dorsal (Do) sides. **(B)** Representative images of pERK in undamaged tissue (top, hair follicles in the inset) and 10 minutes after punch (bottom, pERK^+^ cells at the injury tip in the inset). **(C)** Zoom-in of punched area in *Acomys* ear. Indicated cell subsets activating ERK at 10 minutes after injury: epidermis (Ep), cartilage (Ca), perichondrium (Pe), muscle (Mu), vasculature (Va) labeled by alpha-smooth muscle actin (α-SMA, white); arrowheads indicate double positive cells. **(D)** Quantification of ERK activation at 10min after punch over day 0 (uninjured) within each species. **(E** and **F)** Representative images of cytoplasmic and nuclear pERK in the proximal ear segment at 1hpi (E) and 24hpi (F), insets in the bottom panels, arrowheads indicate pERK^+^ cells at the injury tip. **(G)** Quantification of pERK^+^ nuclei over total nuclei in different skin compartments at the injury site, during the early phase of wound response (P values left to right: 0.0863, 0.0004, 0.0005; 0.0002, <0.0001, 0.0003; 0.0043, 0.0005, 0.0376; 0.2248, 0.0174, 0.0009; 0.9858, 0.0609, 0.1166). Scale bars: 20um. Two-tailed Mann-Whitney test (D), two-way RM ANOVA with Sidak’s multiple comparisons test (G), n=3-4/species/time point. Data are represented as mean ± s.e.m; ns not significant, * P< 0.05, ** P<0.01, *** P<0.001, **** P<0.0001.

### Sustained ERK activation after injury is a hallmark of tissue regeneration

Following the initial burst of ERK activity in response to injury, we evaluated ERK activation during the late phase of wound resolution when the injury response transitions to fibrotic repair in *Mus* or regeneration in *Acomys* (*4*). In *Mus*, ERK activation significantly decreased over time and we observed very few scattered pERK^+^ cells throughout the fibrotic tissue in the distal (fig. S2, A to C) and proximal ear segments (Fig. 2, A to C, and fig. S2, D to G). We observed a similar decrease in *Acomys*, but only in the distal ear segment from D5 to D25 (days post-injury) (fig. S2, A to C). Conversely, in *Acomys* we found persistent and robust ERK activation in the proximal ear segment during wound closure and later during regeneration (Fig. 2, A to D, and fig. S2, D to G). This was consistent with previous observations of asymmetric regeneration in *Acomys* where new tissue formation occurs preferentially in the proximal ear segment, whereas during fibrotic repair, scar tissue is produced equally around the circular punch (*4, 5*). Additionally, pERK^+^ keratinocytes were found at the epidermal margins during wound closure in *Acomys* at D10, consistent with the need for the new epidermis covering the wound to expand during tissue ingrowth (fig. S2H).

**Fig. 2.**
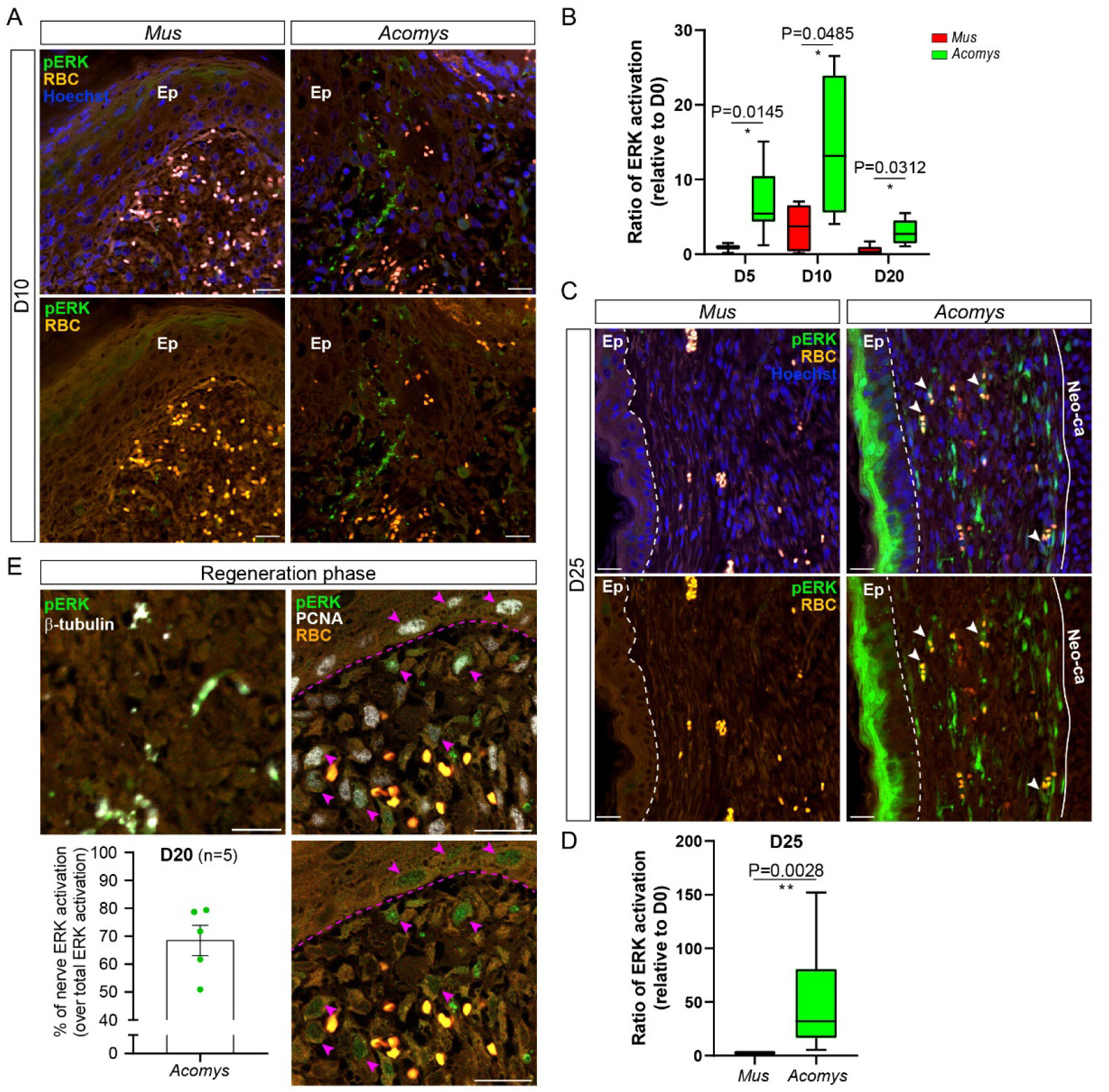
ERK activity is sustained over time only at regenerating wounds. **(A)** Representative images of ERK activation (pERK, green) in the injury tip at D10. **(B)** Box-and-whisker plot of ERK activation levels: bottom whisker corresponds to minimum, box bottom to 25^th^ percentile, box middle to median, box top to 75^th^ percentile, top whisker to maximum. **(C)** Representative images of ERK activation at late time point after ear punch. pERK^+^ keratinocytes in the basal and upper layers of the epidermis (Ep), vasculature (arrowheads, Red Blood Cells RBC, orange), papillary dermal fibroblasts (immediately beneath the epidermis) and cells lining the neo-cartilage in the growing blastema of *Acomys* only (C, right). **(D)** Quantification of pERK levels in *Mus* and *Acomys* at the late phase of wound resolution (n=4 *Mus*, 9 *Acomys*). **(E)** Representative images of pERK^+^ nerves (labeled by β-III Tubulin, white, left) and PCNA^+^ nuclei (white, right) in the regenerating *Acomys* ear at the blastema tip. Arrowheads indicate pERK/PCNA double positive cycling cells; dashes lines demarcates epidermis-dermis boundaries. Quantification of pERK^+^/β-III tubulin^+^ area over the total pERK^+^ area at the injury tip. Scale bars: 20um. Two-tailed unpaired t-test between *Mus* and *Acomys* at each time point. Welch’s t-test correction applied when the variances were unequal (n=3 and 7 for D5, n=5 and 4 for D10, n=6 and 5 for D20 of *Mus* and *Acomys*, respectively) (B), two-tailed Mann-Whitney test (D). Data are represented as mean ± s.e.m (D and E); * P< 0.05, ** P<0.01.

As new tissue began to fill the open hole in *Acomys*, we observed activated ERK throughout the epidermal and mesenchymal connective tissue (blastema). These included keratinocytes, papillary dermal fibroblasts, endothelial cells, mesenchymal cells adjacent to the regenerating cartilage, newly forming chondrocytes (Fig. 2C and fig. S2G) and nerves labeled by β-III tubulin (Fig. 2E, left). At D20, approximately 70% of the pERK signal was detected in nerves at the blastema tip (Fig. 2E). Conversely, pERK and β-III tubulin^+^ nerves were nearly absent in the distal ear segments (fig. S2I). Notably, ERK was activated by cycling mesenchymal cells and basal keratinocytes, labeled by the cell proliferation marker PCNA, at the blastema edge (Fig. 2E, right). Together, these data demonstrate that tissue damage initiates a generic burst of ERK activation in the earliest stages of wound healing. However, robust ERK activation is maintained only in *Acomys* over time and only in the proximal ear segment concurrent with blastema formation and regeneration.

### ERK activation is required independently for wound closure and regeneration

To further investigate the potential role of ERK activation for re-epithelialization and regeneration, we treated the spiny mice with the highly selective small molecule inhibitor Mirdametinib (PD0325901, hereon referred as PD) that prevents ERK phosphorylation and, thereby, its activation (*12*). In contrast to the strong ERK activation observed in the punched ears of the *Acomys* control group, the phosphorylation of both ERK1/2 isoforms (P-p44/P-p42) was strongly reduced upon PD treatment in the epidermis and dermis at 6hpi (Fig. 3, A and B). A series of inhibition experiments was designed to inhibit the early and late phase of ERK activation separately or simultaneously (Fig. 3C). Re-epithelialization occurred in the control group by D12 (Fig. 3D) with new tissue growth closing the ear hole by D40 (Fig. 3, D and E). In contrast, continuous ERK inhibition blocked regeneration and we observed incomplete re-epithelialization and a persistent scab until at least D20 (Fig. 3, D and E). Similarly, animals treated with the early PD scheme exhibited severely delayed wound closure and impaired regeneration (Fig. 3, D and F).

**Fig. 3.**
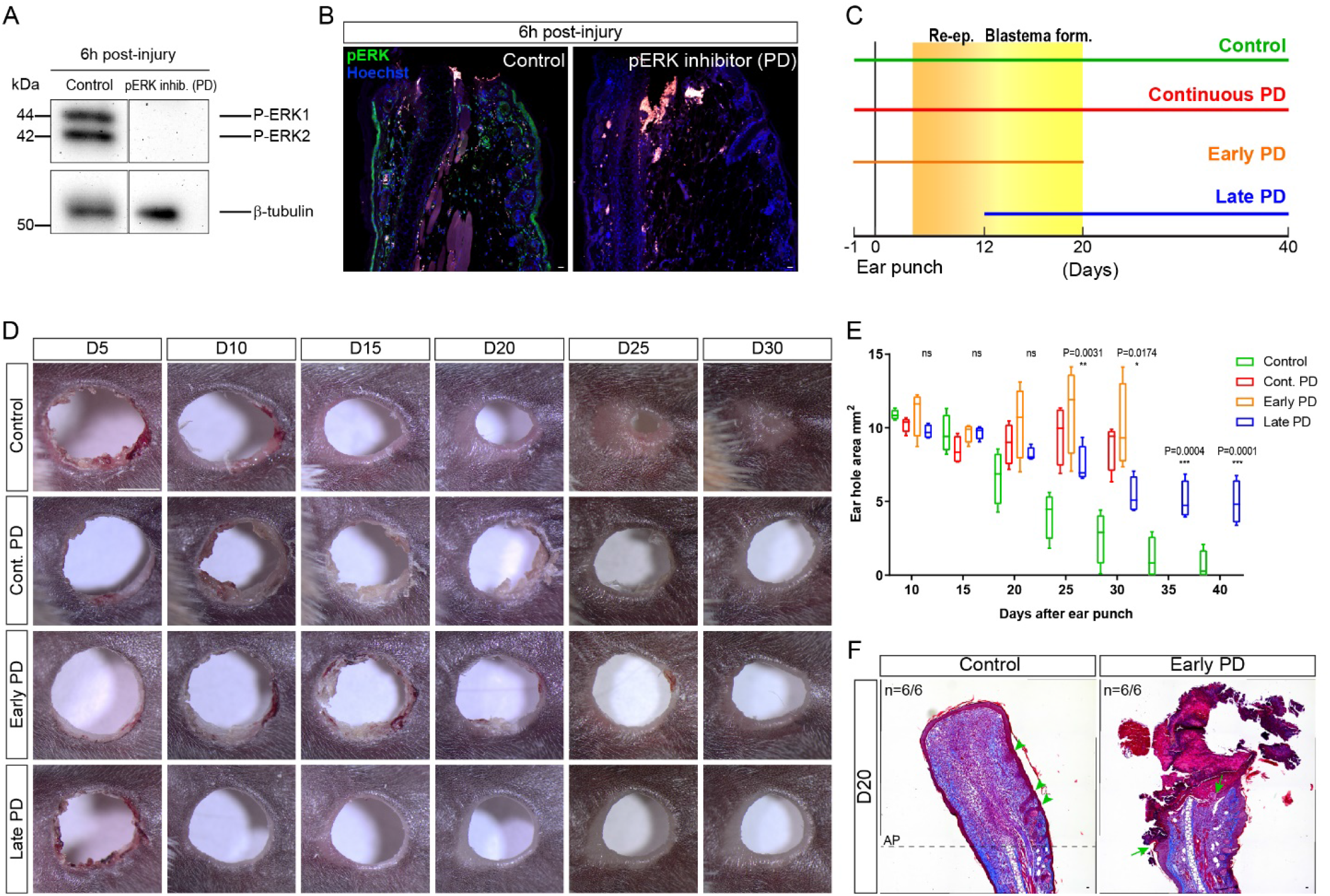
Wound closure and regeneration are independently impaired upon ERK inhibition in *Acomys*. **(A** and **B)** Pharmacological inhibition of ERK activation by Mirdametinib (PD) treatment, confirmed by Western Blot (P-p44/P-p42, phosphorylated ERK1/2 isoforms; β-tubulin as loading control n=3/group) (A) and immunofluorescence (pERK, green, n=3/group) (B) at 6h post-injury. **(C)** Experimental design of ERK inhibition in *Acomys*. PD treatment started after re-epithelialization (Re-ep) at D12 in the Late PD group. **(D)** Representative images of ear hole closure of the same *Acomys* ear after 4-mm punch. Proximal to distal ear segment, left to right, respectively. **(E)**, Box-and-whisker plot of ear hole closure: bottom whisker corresponds to minimum, box bottom to 25^th^ percentile, box middle to median, box top to 75^th^ percentile, top whisker to maximum (n=4/group). Two-way RM ANOVA with Sidak’s multiple comparisons test. **(F)** Representative images of Masson’s Trichrome staining showing severe impairment of wound closure in the Early PD group. New tissue growth and *de novo* appendage formation (indicated by arrowheads) in control ears (left, AP amputation plane), open wound with scab formation in Early PD group (arrows indicate the two wound margins, right). Scale bars: 20um (B and F), 2mm (D); ns not significant, * P< 0.05, ** P<0.01, *** P<0.001.

Because wound closure is necessary for regeneration (*13, 14*), we specifically targeted the late phase of ERK activation associated with new tissue formation by starting the PD treatment at D12 after re-epithelialization occurred (Fig. 3C). Spiny mice undergoing late PD treatment completed re-epithelialization normally, but regeneration was still markedly compromised compared to the control group, supporting a specific role for the sustained ERK activity during regeneration (Fig. 3, D and E). These data demonstrated that ERK activation is necessary for wound closure and its sustained activity is critical for regeneration.

### ERK inhibition shifts the *Acomys* regeneration towards fibrotic repair

To characterize the ERK-dependent molecular events driving blastema formation and regeneration in *Acomys*, we selectively inhibited ERK activation after re-epithelialization (D12) and performed RNA-seq analyses on the proximal ear segments collected at D15, after only 72h of PD treatment (fig. S3, A and B). The efficiency of the PD was confirmed immunohistochemically at D15 (fig. S3, C and D). RNA-seq analysis comparing PD-treated to control punched ear tissue revealed 265 differentially expressed genes (False Discovery Rate, FDR<0.1), 120 of which could be categorized into processes associated with blastema formation and expansion (Fig. 4A). These processes included gene sets associated with cell proliferation, immature wound epidermis (specialized epithelium of activated keratinocytes), production of a pro-regenerative extracellular matrix (ECM), resolution of excessive inflammation and nerve re-growth (*13*).

**Fig. 4.**
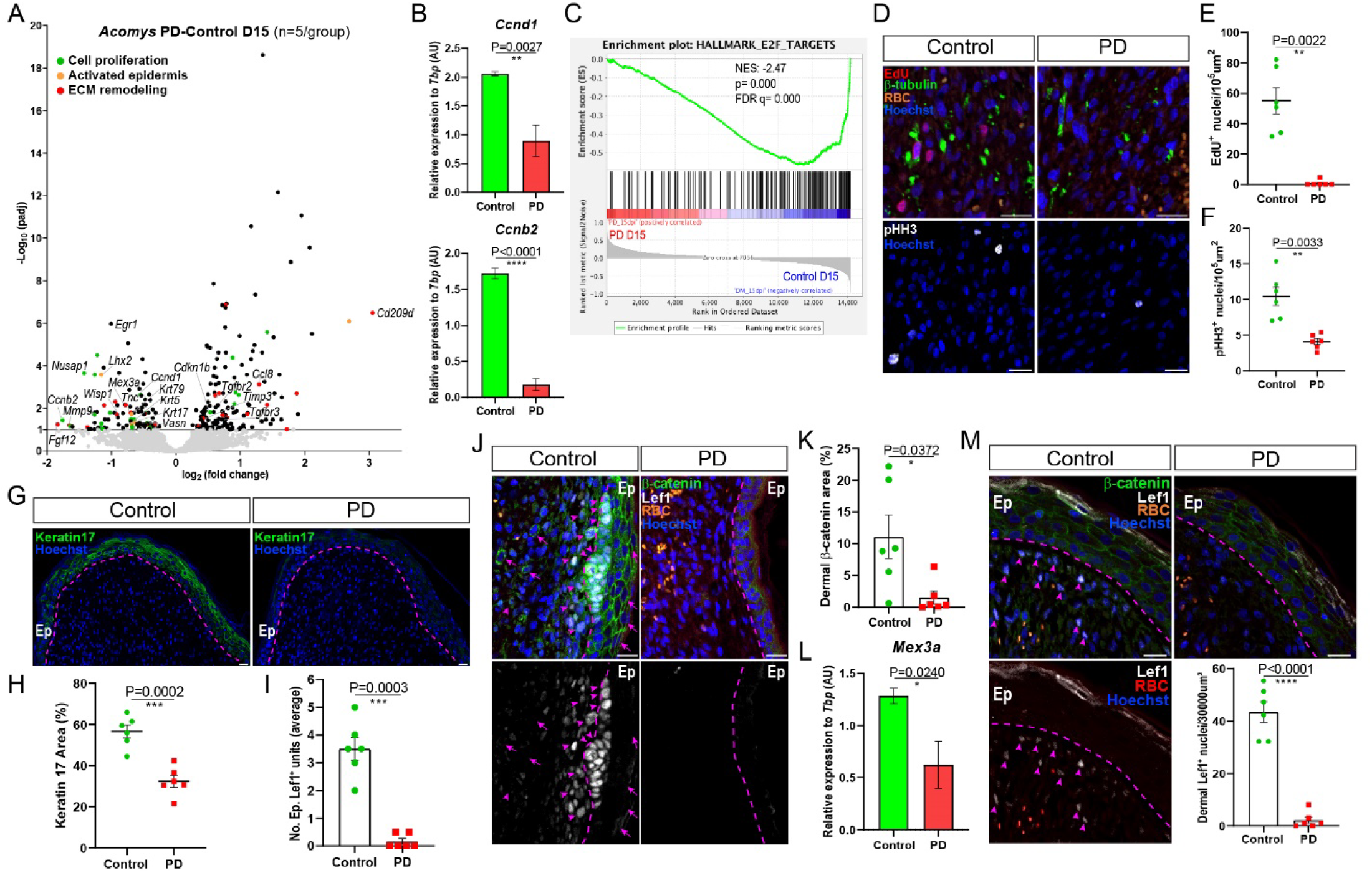
ERK inhibition shifts skin regeneration towards a scar-prone wound response. **(A)** Volcano plot displaying DEGs. Color-coded categories. **(B)** RT-qPCR validation (n=5/group). **(C)** GSEA enrichment plot showing a drastic downregulation of E2F targets gene set in the PD group compared to control at D15. Normalized Enrichment Score (NES), False Discovery Rate (FDR), nominal p-value (p) (n=5/group). (**D** to **F**) Representative images and quantification of EdU^+^ (red, top) and pHH3^+^ (white, bottom) nuclei in the injury area (D15); β-III tubulin (green) (n=6/group). **(G** and **H)** Representative images and quantification of epidermal Keratin 17^+^ (green) area in the injury tip at D15 (n=6/group). (**I**) Quantification of epidermal Lef1^+^ units (n=6/group). (**J**) Representative images of Lef1^+^ nuclei (white) at the hair germ (D15). Cytoplasmic (arrows) and nuclear (arrowheads) β-catenin (green). **(K)** Quantification of dermal β-catenin^+^ area at D15 (n=6/group). **(L)** RT-qPCR validation of *Mex3a* at D15 (n=5/group). **(M)** Representative images and quantification of dermal Lef1^+^ (white) nuclei in the injury tip (D15). Pink dashed line, epidermis (Ep)-dermis boundaries. RBC (red blood cells), nuclei (Hoechst, blue). Scale bars: 20um. Two-tailed unpaired t-test (B, H and L); Two-tailed Mann-Whitney (E). Two-tailed unpaired t-test followed by Welch’s correction (F, I, K and M). Data are represented as mean ± s.e.m; * P< 0.05, ** P<0.01, *** P<0.001, **** P<0.0001.

Consistent with previous work demonstrating a role for ERK signaling in cell cycle progression through the G1/S checkpoint *in vitro* (*8, 15*), our RNA-seq analysis revealed that genes involved in the G1/S transition, including *cyclin D1* (*Ccnd1*) were downregulated after PD treatment *in vivo*. In addition, other genes involved in cell cycle progression were identified; the G2/M-specific *cyclin B2* (*Ccnb2*) and the gene encoding for the mitotic regulator *Nusap1* were downregulated upon ERK inhibition, while suppressors of cell cycle entry, such as *cyclin-dependent kinase inhibitor 1B* (*Cdkn1b, p27*^*Kip1*^), known to promote G1 arrest, were upregulated (Fig. 4, A and B). Gene set enrichment analysis (GSEA) identified the E2F target collection as the top downregulated gene set in the PD-treated *Acomys* compared to the control group at D15 (Fig. 4C). E2F factors are transcriptional regulators of cell cycle genes (*16*) and were markedly downregulated upon ERK inhibition (Fig. 4C). To provide further evidence to the ERK-mediated cell cycle progression during regeneration, we quantified the number of cells actively undergoing S-phase (EdU^+^) and those undergoing mitosis (phospho-histone H3 - pHH3^+^). Cellular analysis revealed a strong reduction of EdU^+^ and pHH3^+^ cells in the proximity of the punched area upon ERK inhibition (Fig. 4, D to F). Analyzing cells across the dorsoventral axis of the ear pinna, we also found that cell density and the number of EdU^+^ cells were higher in the dorsal part of the ear where the majority of hair follicles and nerves are localized (fig. S3, E to G). These data demonstrate that ERK activation is required for sustained cell proliferation *in vivo* during regeneration and suggest that ERK drives active cell cycle progression by modulating the expression of cyclin D1, cyclin B2 and E2F downstream targets.

While the role of ERK was not limited to cell proliferation during regeneration, key hallmarks of blastema formation were also impaired upon ERK inhibition (Fig. 4A). During the transition from wound healing to regeneration, the formation and maintenance of an immature, transitional epidermis (i.e., wound epidermis) is a critical step for blastema formation and expansion of proliferating cells (*4, 17*). The wound epidermis of axolotls and spiny mice retains an embryonic-like organization of keratins, specifically the expression of wound-induced Keratin 17 (*Krt17*) and the dividing basal keratinocyte marker Keratin 5 (*Krt5*) (*3, 4, 17*). Conversely, after a transient, wound-induced increase, *Krt17* expression declines to basal levels as wounds in *Mus* undergo fibrotic repair (*4*). Consistent with the RNA-seq analysis showing that *Krt17* and *Krt5* expression were downregulated in PD-treated samples (Fig. 4A), Keratin 17 protein levels were markedly decreased in keratinocytes overlying the injury upon ERK inhibition in *Acomys* (Fig. 4, G and H, and fig. S3H). These data indicate that sustained ERK activation is necessary to maintain the wound epidermis as a transitional epithelium during blastema expansion.

During complex tissue regeneration in *Acomys*, hair follicle neogenesis occurs from the amputation plane towards the ear punch center as new tissue production closes the open hole (*3*). Our RNA-seq analysis identified another keratin protein, encoded by the *Keratin 79* gene, that was downregulated following ERK inhibition (Fig. 4A). Keratin 79 is expressed by migratory keratinocytes involved in hair follicle morphogenesis (*18*). To assess *de novo* hair follicle formation, we evaluated the presence of the hair placode marker Lef1 and found no Lef1^+^ nuclei in the epidermis of PD-treated *Acomys* compared to the control group (Fig. 4, I and J, and fig. S3I). Lef1 belongs to the T cell Factor (TCF)/LEF family of transcription factors that mediate the nuclear response of Wnt/β-catenin pathway (*19*). Interestingly, nuclear β-catenin co-localized with Lef1^+^ nuclei and β-catenin protein levels decreased upon ERK inhibition (Fig. 4, J and K, and fig. S3, J and K). The inability to generate hair placodes following ERK inhibition was also supported by the downregulation of the LIM homeobox gene *Lhx2* and *Mex3a* in PD-treated *Acomys* (Fig. 4, A and L). The RNA-binding protein Mex3a labels a subpopulation of stem cells promoting regeneration in mouse intestinal epithelium and localizes to hair follicles of the mouse skin (*20, 21*). Lhx2 controls growth and differentiation of hair follicle progenitors in mice (*22*) and progenitor proliferation through Wnt/β-catenin signaling (*23*). We also tested whether ERK inhibition affected Lef1 levels in the dermis. We found mesenchymal cells positive for nuclear Lef1 beneath the hair germ in the papillary dermis (Fig. 4J) and, interestingly, at the blastema tip during regeneration (Fig. 4M). In contrast, Lef1^+^ nuclei were almost entirely absent from the dermis in the PD-treated animals, (Fig. 4, J and M). These data show that Lef1^+^ mesenchymal cells naturally emerge during regeneration in the *Acomys* ear pinna and that the hair follicle neogenesis is ERK-dependent.

The GSEA through the Pathway Interaction Database (PID) (*24*) further supported that genes associated with the nuclear β-catenin signaling were downregulated in the PD-treated group at D15 (fig. S3, L and M). Specifically, core enriched β-catenin pathway-related genes were downregulated upon ERK inhibition, among which *Mmp9* was one of the top differentially expressed genes (fig. S3M). The metalloproteinase Mmp9 is an ECM-degrading enzyme, known to degrade collagen fibers and to promote cell migration during newt limb re-growth after amputation and spiny mouse ear regeneration (*4, 25, 26*). To gain further insights into the ERK-dependent matrisome, we analyzed the top differentially expressed genes between PD and control groups against the murine ECM atlas (http://matrisomeproject.mit.edu) (*27*). In addition to *Mmp9*, we identified other key matrisome genes whose expression was altered upon ERK inhibition. Among those were genes encoding secreted factors (*Fgf12, Ccl8*), ECM regulators (*Cd209d*), ECM-affiliated proteins (*Timp3*) and core matrix proteins (*Wisp1, Tnc*) (Fig. 5, A and B). Interestingly, the gene encoding Timp3, a tissue inhibitor of metalloproteinases that binds and irreversibly inactivates collagenases and metalloproteinases, such as Mmp9, was upregulated in the PD group (Fig. 4A and 5A). Notably, *Fgf12* and pro-regenerative genes, including *Tnc*, the Wnt/β-catenin downstream effector Wnt-1 inducible signaling protein 1 (*Wisp1*) and *Mmp9* were significantly downregulated upon ERK inhibition (Fig. 5, A and B). Wisp1 acts as a secreted matricellular signal regulating ECM production (*28, 29*). In stark contrast, *Cd209d* and pro-fibrotic genes, including *Ccl8 (30)*, were upregulated in the PD group (Fig. 5, A and B). Recent work identified a pro-regenerative ECM profile in *Acomys*, with enrichment for *TnC, Mmp9* and low levels of *Col1a1* (encoding type I collagen) (*4*). Conversely, *Mus* fibrotic repair was characterized by abundant *Col1a1* and reduced *Tnc* and *Mmp9* expression. While we found Mmp9^+^ cells in the blastema and among basal keratinocytes of the wound epidermis, the number of Mmp9^+^ cells significantly decreased upon ERK inhibition (Fig. 5, C and D, and fig. S3N).

**Fig. 5.**
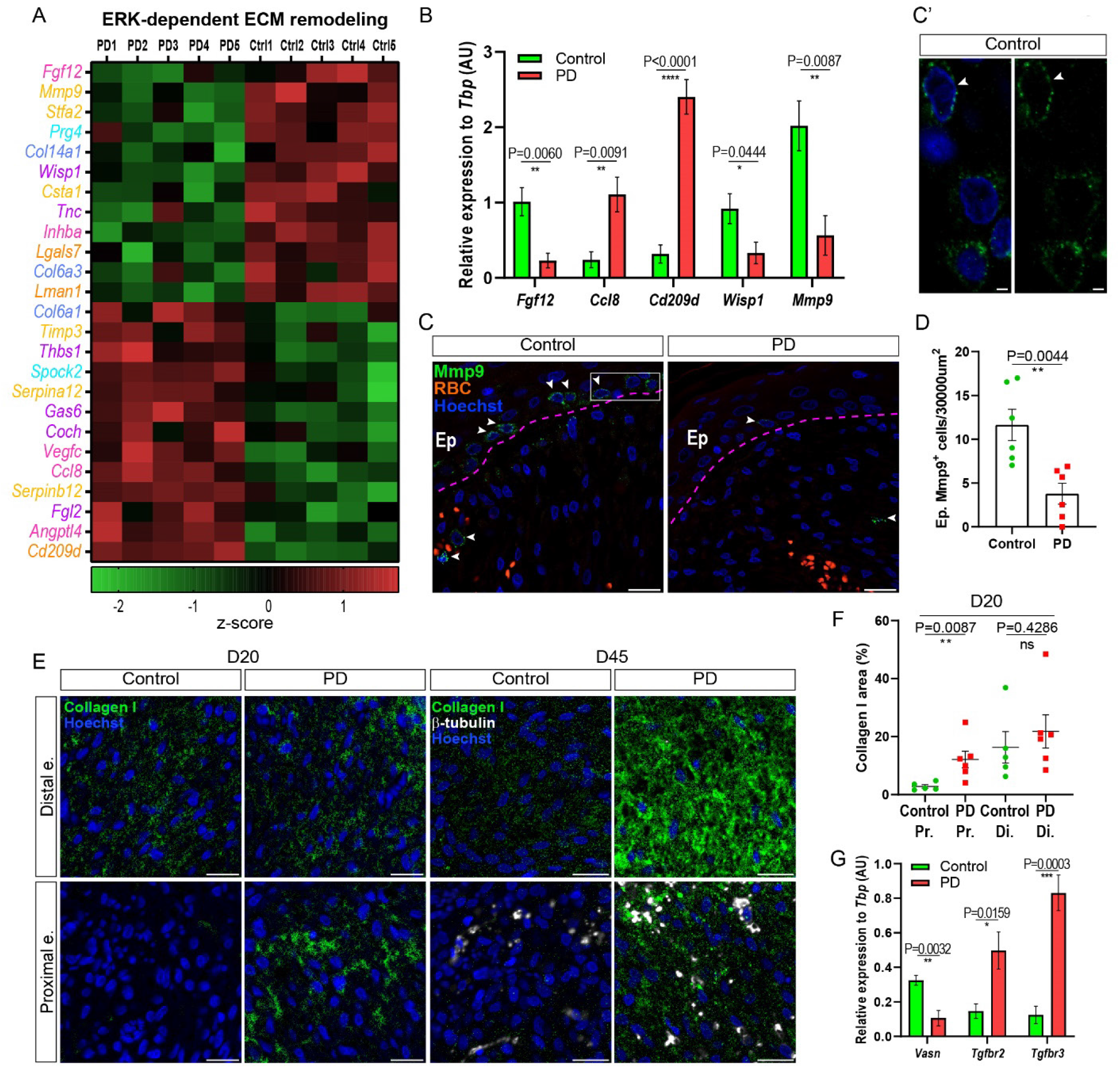
ERK inhibition converts the regeneration-associated matrisome into a pro-fibrotic ECM profile. **(A)** ERK-dependent ECM remodeling (color-coded categories in Methods). **(B)** RT-qPCR validation of matrisome genes (D15, n=5/group). **(C and D)** Representative images and quantification of Mmp9^+^ cells (green) in the injury tip at D15 (n=6/group). Zoom-in of Mmp9^+^ cells in the basal layer of the epidermis (C’). **(E and F)** Representative images and quantification of collagen 1a1 (green) in the injury area of the proximal and distal ear segments at D20 (n=5 control, 6 PD) (left) and D45 (n=4/group) (right); β-III tubulin (white). **(G)** RT-qPCR of TGF-β signaling components (D15, n=5/group). Scale bars: 20um; 2um in inset (C’). Gene expression value normalized by z-score transformation; (n=5/group) (A). Two-tailed unpaired t-test (B, D and G), two-tailed Mann-Whitney (F). Data are represented as mean ± s.e.m; ns not significant, * P< 0.05, ** P<0.01, *** P<0.001, **** P<0.0001.

Due to the collagen-degrading activity of Mmp9 and the close association of type 1 collagen with fibrosis (*31, 32*), we quantified collagen deposition upon ERK inhibition and found significantly higher levels of collagen I in the proximal ear segment of PD-treated spiny mice compared to the control group at D20 and D45 (Fig. 5, E and F, and fig. S3O). Notably, no differences were observed in the distal ear segment of the PD and control group, where cell proliferation was nearly absent (Fig. 5, E and F, and fig. S3O). Among the enriched subsets of the Gene Ontology Cellular Components (GO-CC), the gene sets related to ECM and collagen were found to be significantly enriched upon PD treatment (fig. S3P).

Furthermore, TGF-β signaling is a master regulator of ECM assembly and fibrogenesis and is upregulated in organ fibrosis (*33*). Accordingly, PD-treated spiny mice exhibited significantly higher levels of *Tgfbr2* and *Tgfbr3* expression and reduced levels of vasoneurin (*Vasn*), a negative regulator of TGF-β signaling (*34*) (Fig. 5G). Thus, our data indicate that the inhibition of ERK activation during regenerative healing in *Acomys* shifts the healing trajectory towards fibrotic repair.

### Wound-induced and regeneration-specific ERK targets contain AP-1 binding sites in mammals

To investigate whether ERK activation coordinates a regenerative program that can explain its prominent effect on blastema formation in terms of cell proliferation, the formation of an active wound epidermis and ECM remodeling, we focused our analysis on transcription factor binding sites enriched in ERK-dependent genes. GSEA analysis for canonical pathways identified significant changes in the activating protein-1 (AP-1) transcription factor network, upon ERK inhibition at D15 (Fig. 6A). When we interrogated our RNA-seq dataset against collections of transcription factor targets (*35*), the AP-1 binding site, TGANTCA, was one of the most enriched motifs (fig. S4A).

**Fig. 6.**
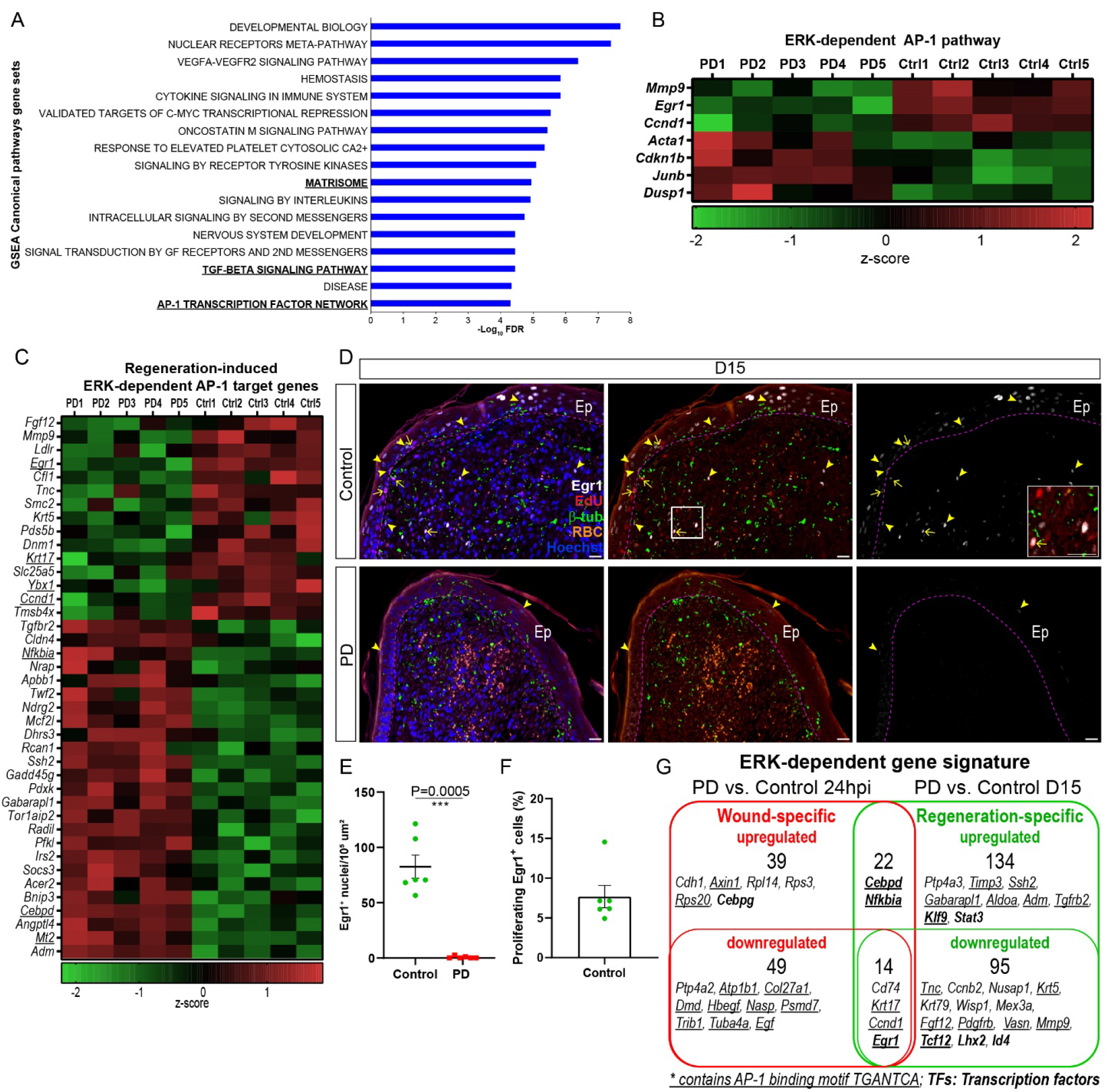
ERK activity modulates AP-1 target genes during blastema formation. **(A)** GSEA Canonical pathway analysis showing statistically significant gene sets (y axis) enriched upon ERK inhibition. X axis indicates -Log_10_ False Discovery Rate (FDR) statistical significance (n=5/group). **(B)** Heatmap showing ERK-dependent genes encoding for components of the AP-1 signaling, among the top DEGs at D15. **(C)** Heatmap showing the top 40 DEGs at D15 containing the AP-1 binding site. Underlined the ERK-dependent DEGs whose expression was wound-induced (at 24h post-injury) and sustained also during the regeneration phase; green and red colors indicate downregulated and upregulated genes, respectively. **(D)** Representative images of ERK downstream target Egr1 (white); β-III tubulin (green), EdU (red), RBC (orange); arrowheads indicate Egr1^+^ cells, arrows indicate Egr1^+^/EdU^+^ cells. Inset showing Egr1^+^/EdU^+^ double positive cells at the blastema tip in the control group. **(E)** Quantification of Egr1^+^ nuclei at D15 (n=6/group). **(F)** Quantification of proliferating Egr1^+^ cells over total Egr1^+^ cells at D15 (n=6). **(G)** Venn diagram showing genes up- and down-regulated in PD at 24hpi and D15. In the intersection, genes induced by injury and sustained during regeneration. Pink dashed line, epidermis (Ep)-dermis boundaries. RBC (red blood cells), nuclei (Hoechst, blue). Scale bars: 20um; Two-tailed unpaired t-test followed by Welch’s correction (E). Data are represented as mean ± s.e.m; *** P<0.001.

AP-1 binding sites can act as responsive elements driving the expression of genes with opposing outcomes, from regeneration (*36*) and proliferation (*37*) to stress response, apoptosis, inflammation (*38*) and fibrosis (*32, 39*). Recently, the AP-1 motif was found enriched in injury-responsive enhancers driving both injury and regenerative responses in fish (*36*) and AP-1 factors induced cell cycle progression by modulating cyclins and E2F, transcriptional regulators of cell cycle genes, *in vitro* (*16*). We found that Cyclin D1, Cyclin B2 and E2F target genes were ERK-dependent *in vivo* (Fig. 4, A to C) and top DEGs identified at D15 were part of the AP-1 transcription factor network (Fig. 6B). Furthermore, 77 of 265 top differentially expressed genes between PD-treated and control group displayed at least one predicted AP-1 binding site in their promoter region (Fig. 6C). Notably, among these ERK-dependent AP-1 target genes, pro-regenerative genes, including *Ccnd1* (*40*), *Keratin 17* (*41*), *Mmp9*, and *Egr1* (*42*) (downregulated in PD-treated *Acomys*) can be differentiated from pro-fibrotic genes, including *Tgfbr2* (upregulated in PD-treated *Acomys*) (Fig. 6C). Interestingly, Egr1 regulates chromatin accessibility and is required for regeneration in acoel flatworms (*43*). We found that *Egr1* expression and the number of Egr1^+^ nuclei were drastically decreased in the ear dermis and epidermis upon ERK inhibition (Fig. 6, C to E). Notably, approximately 8% of the overall Egr1^+^ cells were actively cycling cells, as they were co-labeled with EdU (Fig. 6F). These data suggest that the pro-regenerative effect of AP-1 responsive enhancers depends on the specific genes they regulate and that ERK activation stimulates *Acomys* regeneration at least partially through AP-1 target genes.

In addition, we compared the top differentially expressed genes between control and PD-treated *Acomys* ears isolated at 24h post-injury (injury-specific) with those isolated at D15 (regeneration-specific). This analysis identified an injury-specific and a regeneration-specific ERK gene signature (Fig. 6G). 35 of 124 top differentially expressed genes activated in the early injury response (24h post-injury) contained at least one putative AP-1 binding site in their promoter region (fig. S4B). Interestingly, the expression of some of these injury-induced ERK-dependent genes, including *Keratin17, Ccnd1* and *Egr1*, were later maintained during the regeneration phase, resembling the sustained ERK activation in *Acomys* (Fig. 6C and fig. S4B). Notably, also some regeneration-specific ERK-dependent genes (e.g., *Keratin5, Tnc, Mmp9*) revealed putative AP-1 target sites (Fig. 6G). Together, these data indicate that a regenerative response is coordinated by regeneration-specific ERK target genes in combination with the sustained expression of early wound-induced ERK-dependent genes. ERK activation may therefore be the missing link that connects the injury signals to AP-1 mediated healing outcomes.

### ERK activity correlates with regeneration rate and is regulated by FGF and ErbB2 receptors during mammalian blastema formation

Ear punches in *Acomys* regenerate asymmetrically with new tissue production occurring from the proximal area of the hole (*13*). In line with this, we found that the regeneration rate was higher when deploying the 4-mm ear punch proximal to the head compared to a more distally placed punch in the other ear of the same spiny mouse (Fig. 7, A and B). Notably, proximal ear punches regenerated completely within 20 days (95.58% ± 3.13 of punch area), while the hole of distal punches was filled with new tissue by one third only (26.75% ± 3.74 of punch area) at D20 (Fig. 7, A and B). The regeneration rate of the proximal ear punch was 215 ± 34 um/day between D10 and D15, almost three times faster than a distal ear punch (70 ± 14 um/day) (Fig. 7C).

**Fig. 7.**
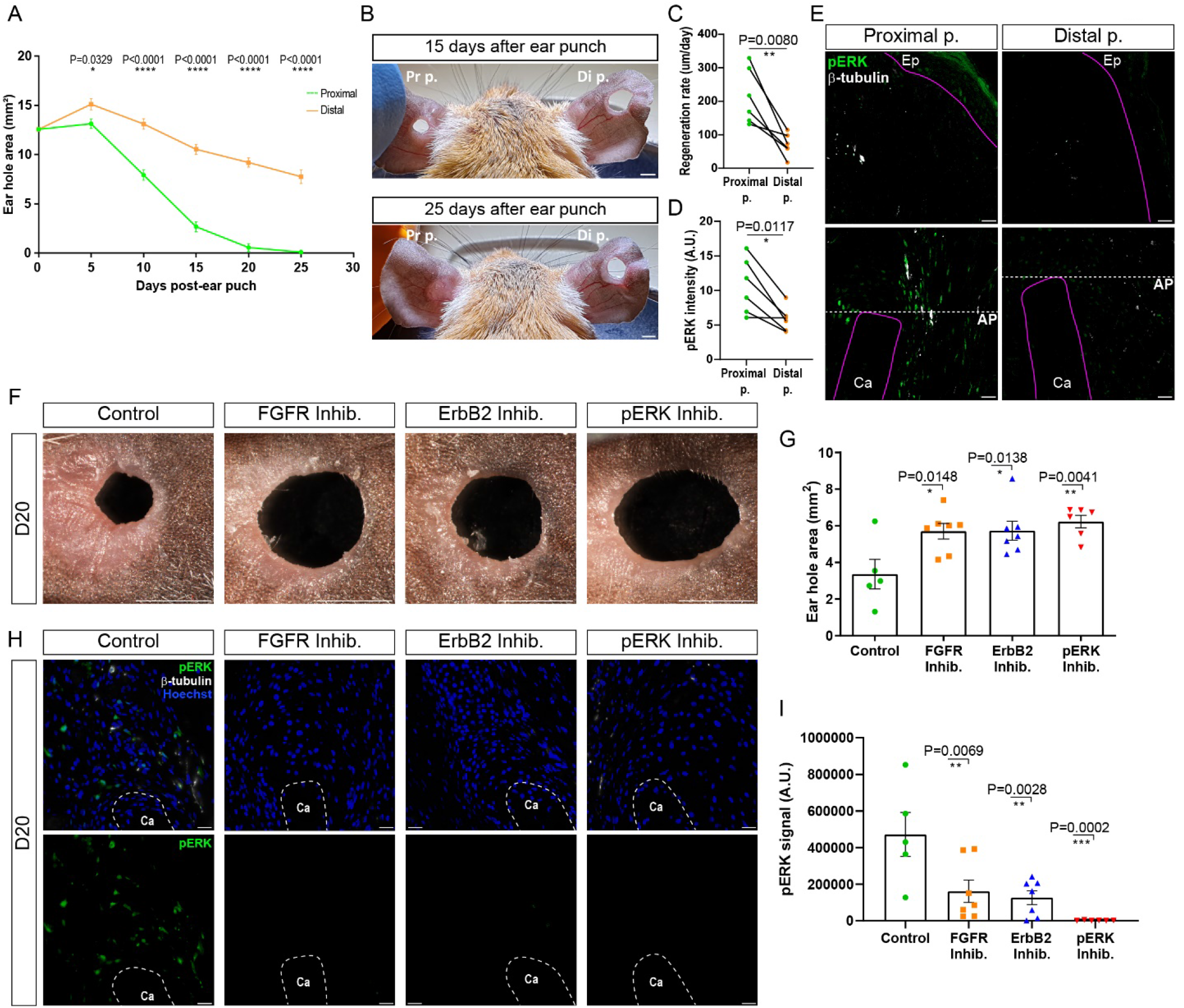
ERK activity correlates with regeneration rate and is regulated by FGFR and ErbB2 receptors during regeneration. **(A)** Ear hole closure in proximal and distal punch (n=6). **(B)** Representative images of hole closure after proximal (left) and distal (right) 4-mm punch. **(C)** Paired analysis of regeneration rate between proximal and distal punches. **(D)** Paired analysis of ERK activation: lines connect proximal to distal punch in the same mouse (n=6). **(E)** Representative images of pERK (green, arrowheads) at injury tip (top) and amputation plane (AP) (bottom), β-III tubulin (white). (**F** and **G**) Representative images and quantification of ear hole closure after inhibition of FGFR (n=7), ErbB2 (n=7) or pERK (n=6) compared to the control group (n=5) at D20. (**H** and **I**) Representative images and quantification of ERK activation (pERK, green) in control (n=5), FGFR (n=7), ErbB2 (n=7) or pERK (n=6) inhibitor groups; β-III tubulin, white, intact cartilage (Ca) of the pre-amputation plane. Scale bar: 2mm (B and F), 20um (E and H). Two-way RM ANOVA with Sidak’s multiple comparisons test (A), two-tailed paired t-test (C and D), one-way ANOVA with Dunnett’s multiple comparisons test (G and I). Data are represented as mean ± s.e.m; * P< 0.05, ** P<0.01, *** P<0.001, **** P<0.0001.

As ERK activity is required for *Acomys* regeneration, we asked whether the faster regeneration rate correlated with higher ERK activity. We found significantly higher pERK levels in proximal compared to distal injuries, which positively correlated with ear closure rate (Fig. 7, D and E, and fig. S5A). These data indicate that the extent of ERK activity is a key determinant for tissue growth and regeneration and suggest that exogenous factors delivered to the injury via peripheral nerves might control ERK signaling.

Since nerves are necessary for blastema expansion and are crucial sources of the mitogens fibroblast growth factors (FGFs) and neuregulins (*44, 45*), we sought to investigate whether antagonizing receptors for these mitogens could inhibit regeneration in *Acomys*. By pharmacologically inhibiting the FGFR and Erbb2 (Erb-B2 Receptor Tyrosine Kinase-2) receptors from D12 (after re-epithelization following 4mm ear punch) to D20, we found that regeneration and ear punch closure were strongly impaired compared to the control group (One-way ANOVA, F=5,179 P=0.0078) (Fig. 7, F and G). Notably, ERK activity significantly decreased by 66% and 73% in the FGFR inhibitor and ErbB2 inhibitor groups, respectively (One-way ANOVA, F=8.816, P=0.0006) (Fig. 7, H and I). These data indicate that FGFR and ErbB2 act as upstream ERK regulators during composite tissue regeneration in spiny mice.

### ERK activators stimulate a regenerative response in scarring wounds

To test whether regeneration could be induced in normally non-regenerating *Mus* wounds, we ectopically released recombinant FGF2 and Neuregulin-1 (Nrg1) in the *Mus* ear pinna following a 4-mm ear punch, that identifies regenerative abilities in mammals (*4*). We locally implanted micro-beads soaked with FGF2 and Nrg1 after re-epithelialization occurred, at D10 (Fig. 8A). We found that a cocktail of FGF2/Nrg1 led to a significant increase in tissue growth compared to the control group at D20 (Fig. 8, B and C). To confirm that FGF2/Nrg1 enhanced ERK activation, we assessed ERK activity and EdU incorporation at D13 and D20. In contrast to the PBS-soaked beads, the ears implanted with FGF2/Nrg1 beads showed significantly enhanced ERK activity and an increased number of EdU^+^ cells in the epidermis and dermis (Fig. 8, D to G, and fig. S5B and C). This manifested into a higher cell density around the FGF2/Nrg1-soaked beads (Fig. 8H). Strikingly, we also observed hair follicle neogenesis at the injury tip in *Mus* wounds at D20, demonstrating that the increased ear hole closure occurred through regenerative healing and not simply through increased scar tissue production (Fig. 8, H and I).

**Fig. 8.**
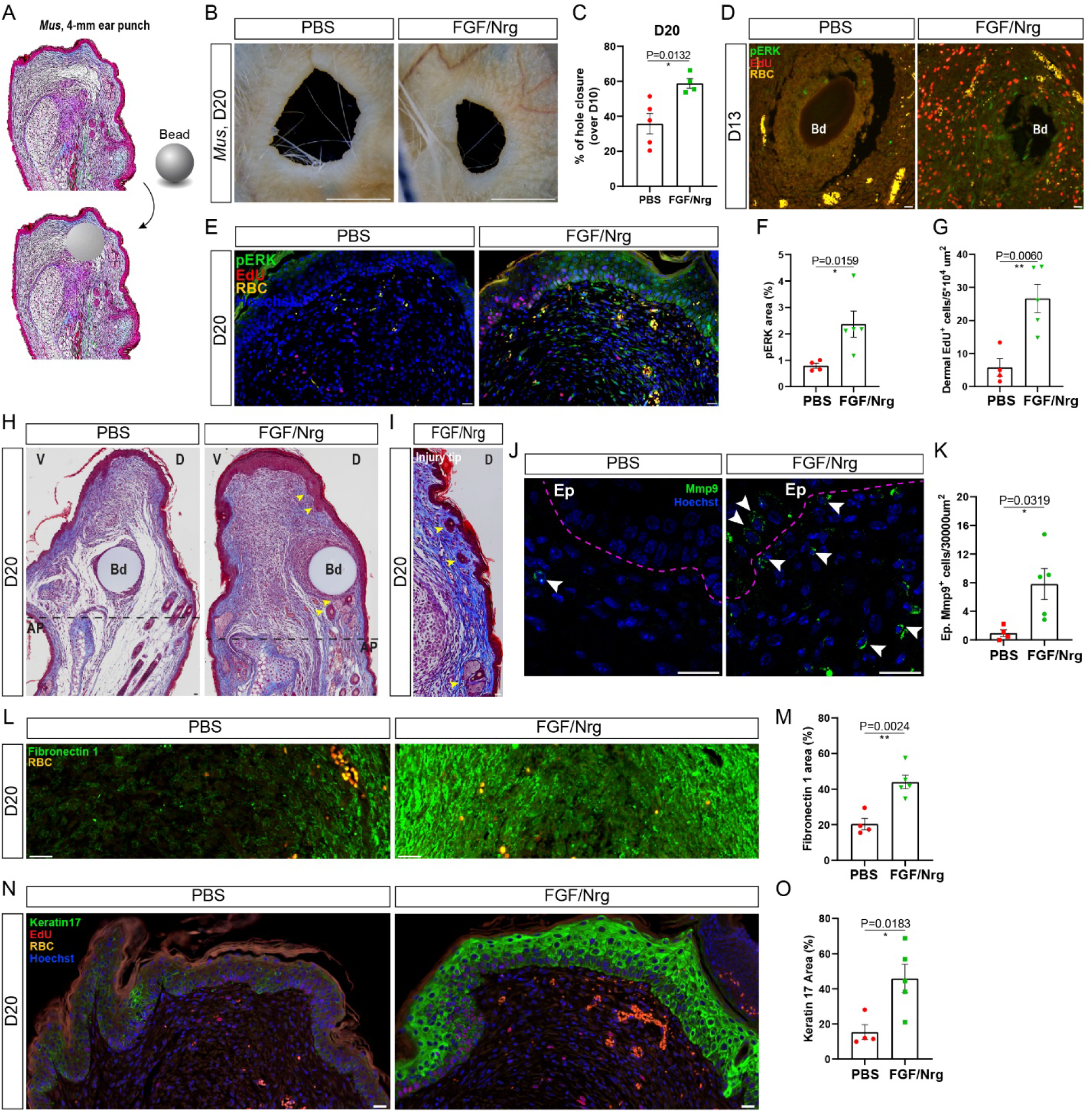
ERK activators induce a regenerative response and new hair follicle formation in normally non-regenerating *Mus* wounds. **(A)** Graphics of the experimental design. Shown a punched ear before and after bead implantation at the injury tip. **(B** and **C)** Representative images and quantification of hole closure after 4-mm ear punch. **(D** and **E)** Representative images of ERK activation and proliferating cells in the group locally treated with FGF/Nrg-soaked beads compared to PBS control. **(F** and **G)** Quantification of ERK activation and dermal proliferating cells in the injury tip at D20 ((n=4 PBS, n=5 FGF/Nrg). **(H)** Representative Masson’s Trichrome staining of the beads-implanted ears; injury tip at the top, bead (Bd), ventral (V), dorsal (D) (n=4 PBS, n=5 FGF/Nrg). **(I)** Representative images of newly formed appendages (arrowheads indicate *de novo* hair follicle formation) at the injury tip (n=5 FGF/Nrg). **(J** to **O)** Representative images and quantification of Mmp9^+^ cells (green, arrowheads) (J and K), Fibronectin 1 (green) (L and M) and Keratin 17 (green) (N and O) at the injury tip at D20 (n=4 PBS, n=5 FGF/Nrg). Pink dashed line, epidermis (Ep)-dermis boundaries. RBC (Red Blood Cells, orange), nuclei (Hoechst, blue); black dashed line, amputation plane (AP). Scale bar: 20um, except 2mm (B). Two-tailed unpaired t-test (C, G, M, O), two-tailed Mann-Whitney test (F), two-tailed unpaired t-test followed by Welch’s correction (K). Data are represented as mean ±s.e.m; * P< 0.05, ** P<0.01.

To test whether the FGF2/Nrg1-soaked beads induced a pro-regenerative microenvironment similar to what observed in spiny mice, we assessed the abundance of Mmp9 and the blastema-associated ECM protein fibronectin 1 (*4, 46*). Notably, the FGF2/Nrg1 mix stimulated a significant increase of Mmp9^+^ cells at the injury tip (Fig. 8, J and K). Moreover, we detected augmented fibronectin 1 levels in the newly produced mesenchyme compared to the PBS control group (Fig. 8, L and M, and fig. S5D).

Furthermore, we analyzed Keratin 17 levels in the epidermis, where its sustained expression is associated with blastemal growth and regeneration, while it is reduced in fibrosis (*3, 17, 47*). Consistent with the enhanced regenerative phenotype observed in *Mus* ears after bead transplantation, FGF2/Nrg1-soaked beads led to the activation and maintenance of Keratin 17 expression in the wound epidermis at D20 (Fig. 8, N and O, and fig. S5E). Keratin 17 levels showed a strong positive correlation with keratinocyte proliferation and ERK activity (fig. S5, F and G). In turn, ERK activation positively correlated with cell proliferation in the dermis of the injury tip (fig. S5H). Thus, the FGF2/Nrg1 mix triggered ERK activation and cell proliferation in the dermis, promoted a fibronectin 1- and Mmp9-enriched microenvironment and prevented differentiation of the epidermis by inducing cell proliferation and Keratin 17 expression in the *Mus* wound epidermis. Together, our data demonstrate that ERK activity is capable of shifting a fibrotic repair program to exhibit hallmarks of regeneration in *Mus* wounds.

## Discussion

The mechanistic basis for naturally occurring complex tissue regeneration in adult mammals remains incompletely understood. Here we uncovered that ERK acts as a master regulator and molecular switch between tissue regeneration and fibrosis in the highly regenerative spiny mouse, a rare example of complex tissue regeneration in an adult mammal. Our data show that ERK activation is required for the wound healing phase of tissue repair and, specifically, during regeneration, where it acts (1) to promote a sustained proliferative response by controlling cyclins and E2F factors, (2) to maintain the wound epidermis as a transitional epithelium during blastema formation, (3) to stimulate a pro-regenerative extracellular matrix remodeling, (4) to activate Lef1-dependent hair follicle neogenesis and (5) to transduce the FGFR and ErbB2 receptors downstream signals during skin regeneration.

While ERK is induced immediately in response to injury regardless of the healing outcome, its activation is maintained only where a blastema forms. Our study demonstrates a functional requirement for ERK activity as ERK inhibition in *Acomys* wounds resulted in a scar-prone phenotype, characterized by pro-fibrotic ECM profile, decreased cell proliferation, reduced expression of Keratin 17 in the epidermis and absence of hair follicle neogenesis. Importantly, we could harness appropriate ERK activity to promote a pro-regenerative environment, increased cell proliferation and tissue regeneration in normally non-regenerative *Mus* wounds.

Our analysis of ERK-dependent gene expression during regeneration in *Acomys* supports these data and suggests that sustained ERK activity after re-epithelialization is required to antagonize fibrosis by maintaining the expression of injury-responsive genes that stimulate a pro-regenerative extracellular environment. Although ERK was immediately activated after injury in *Mus* and *Acomys* wounds, ERK activity was significantly higher and broader in *Acomys* cells. This is consistent with the observation that *Acomys* fibroblasts possess a more euchromatic state compared to the DNA of *Mus* cells *(4)*, possibly resulting in a faster adaptation to extracellular stressors. Previous studies have shown that axolotls retain juvenile traits during adulthood by maintaining high levels of euchromatin which keeps chromatin accessible for a prompt induction of gene expression in response to injury (*48*). Similar to quiescent stem cells, some *Acomys* cells may be retained in an alert state (*49*), allowing them to promptly react to microenvironmental changes. Consistent with this hypothesis, ERK2 has been shown to bind developmental genes in embryonic stem cells and phosphorylate RNA polymerase II which induces a poised state on gene promoters (*50*). Moreover, ERK phosphorylation can lead to chromatin accessibility through histone modifications (*51*) and the transcription of immediate early genes that are highly responsive to injury signals, including AP-1 target genes and *Egr1* which occupies the promoter of the regeneration-responsive enhancers in acoel flatworms (*43*). Similar to the observed ERK activation profile in acoels, AP-1 transcription factors are induced immediately after injury, regulate cell cycle through cyclin D1 (*16*) and promote zebrafish heart regeneration through cardiomyocyte dedifferentiation and proliferation (*52*). Interestingly, we found multiple ERK-dependent genes containing AP-1 binding sites in their promoter region, including the cell cycle gene *Ccnd1* and pro-regenerative genes such as *Egr1* and *Krt17*, suggesting that ERK-driven regeneration is at least partially elicited by specific AP-1 induced injury-responsive genes in mammals. The persistent expression of a multitude of ERK-dependent genes is consistent with ERK regulating the expression of a cellular regeneration program. The expression of ERK targets highlights that regeneration-specific effectors (i.e., Tenascin C, Keratin 5, Mmp9), together with the extent and the duration of wound signals (pERK, Egr1, Keratin 17) play a critical role towards orchestrating a regenerative response.

The ectopic application of ERK activators, FGF2/Nrg1, was sufficient to stimulate *Acomys* regenerative traits in *Mus* wounds, including the ability to induce *de novo* hair follicle formation. While the ERK-dependent mechanisms induced by injury might differ to some extent across species, the role of ERK as a major player at the crossroad between tissue regeneration and fibrosis seems to be conserved across evolution and, importantly, in mammals. It is likely that this is a generally valid concept as even adult *Mus* that normally do not regenerate composite tissues, showed signs of regeneration after ERK was ectopically activated.

While the components of ERK signaling pathway are evolutionarily conserved, the degree to which instances of tissue regeneration represent phylogenetic inertia or *de novo* episodes of convergent evolution remains unknown. Recent findings have revisited the hypothesis that there exists a “regeneration program” which is suppressed by inhibitory mechanisms promoting cell commitment and differentiation to safeguard tissue stability and DNA integrity (*53, 54*). Natural selection may have favored mechanisms for fast wound healing and a strong fibrotic response to protect functional parenchyma at the expense of the plasticity required for regeneration. Thus, the ability to regenerate tissues and organs may not be completely lost, but dormant in the cells of poorly-regenerating mammals, such as adult laboratory mice and humans. Pro-regenerative core signaling pathways, such as MAPK/ERK signaling, are activated in *Mus* cells but their activity appears to be limited, thereby tapering off the transcription of pro-regenerative genes. This supports that fibrotic repair is present because a regenerative response is not sustained. However, extrinsic signals may trigger the latent cell potential, circumvent regeneration blocks and rescue pro-regenerative traits. The presence of dormant regenerative traits in mammals harbors enormous implications for regenerative medicine. Indeed, the identification of specific factors that activate core signaling pathways may awaken a latent regenerative program in mammalian cells. Ultimately, our data propose a model in which ERK activity might be therapeutically fine-tuned to antagonize fibrosis and stimulate regeneration following tissue and organ damage.

## Materials and Methods

### Animals

Spiny mice (*Acomys cahirinus*) and CD1 mice (*Mus musculus*) were housed at the University of Kentucky, Lexington, USA and at the Hubrecht Institute, Utrecht, NL, as previously described (*4*). *Acomys* were fed a 3:1 mixture by volume of 14% protein mouse chow and black-oil sunflower seeds. CD1 mice were fed mouse chow only. *Acomys* and *Mus* experienced a 12:12 (light: dark) light cycle. All the employed animals were sexually mature and age- and sex-matched within experiments. In case of dead or sick animals, the samples were excluded from the study. All animal procedures were approved by the University of Kentucky Institutional Animal Care and Use Committee (IACUC) under protocol 2019–3254 and by the Animal Ethics Committee of the Royal Netherlands Academy of Arts and Sciences (KNAW) (project license AVD8010020187144).

### Surgery and 4-mm ear punch assay

Upon 4% (v/v) vaporized isoflurane-induced anesthesia, a through-and-through hole was performed with a 4-mm biopsy punch in the center of both the left and right ear pinna, throughout the study, except for the proximal vs. distal ear punch experiment described below. At specified time points, the animals were anesthetized and the ear hole area was calculated with a digital caliper, as previously described (*4*). Briefly, orthogonal diameters along the y and x axes were measured, the respective radii, r_y_ and r_x_ were calculated and the ear hole area (A) was obtained: A= πr_y_r_x_.

To detect differences in the ear hole closure rate between a proximal and a distal 4-mm ear punch, the proximal ear punch was performed at approximately 1-mm from the skull of the left ear and the distal ear punch at approximately 4-mm from the tip of the right ear along the y axis, both medially in the ear pinna of the same spiny mouse, as an internal control.

### *In vivo* treatments

After 4mm ear punch, the animals were randomized and subjected to intraperitoneal injections of ERK inhibitor, FGFR inhibitor, ErbB2 inhibitor or vehicle. Pharmacological ERK inhibition was achieved by injecting the highly selective MEK inhibitor Mirdametinib (PD0325901, indicated as PD, purchased by Selleck Chemicals). PD was injected at 18mg/kg in 200ul of total volume per mouse every other day. The Continuous PD group was treated from day -1 across the duration of the experiment. The Early PD group was treated from day -1 to D20. The Late PD group was treated from D12 till the end of the experiment.

For the RNA-seq experiment and RNA-seq data validation, PD was injected daily from D12 to D15 only, at the same dose and formulation.

The selective AZD4547 (*55*) and Mubritinib (TAK165) (*56*) small molecules were used to pharmacologically inhibit the FGFR and ErbB2 receptors, respectively. They were injected as the ERK inhibitor in 200ul of total volume per mouse daily according to manufacturer’s instructions, from D12, after re-epithelialization, to D20 (eight days only).

All the inhibitors were dissolved in a mix of DMSO, polyethylene glycol (PEG), PBS/Tween upon injection, based on manufacturer’s recommendations. The control group was injected with the same mix, except of the inhibitor.

For the rescue studies in *Mus*, after 4-mm ear punch, the animals were randomized and subjected to implantation of micro-beads (BioRad) soaked with ERK activators: FGF2 (Sigma, No. 341583)/Nrg1 (Peprotech, No. 100-03) (dissolved in PBS and evenly mixed). PBS-soaked beads were used as control. Two soaked micro-beads were implanted in the ear punch area after re-epithelialization occurred, at D10 and the ear tissue was collected at D13 or D20 for immunohistochemistry. In the same way, two micro-beads were implanted at D10 and D15 and the tissue was collected at D20 for the assessment of ear closure and hair follicle neogenesis. The respective PBS control group underwent the same procedure.

In order to assess the number of cells in S-phase in *Acomys* and *Mus* wounds, 2mM EdU diluted in 200 ul of saline solution were intraperitoneally injected through three EdU pulses evenly distributed over a period of nine hours before tissue collection. EdU-incorporating cells were detected on formalin-fixed, paraffin embedded tissue sections through copper-catalyzed click reaction by using Alexa Fluor 594 Azide (ThermoFisher Scientific).

### Histology and immunohistochemistry

Upon tissue collection, ear samples were fixed overnight in 10% (v/v) neutral buffered formalin at 4°C, washed in PBS, followed by dehydration and paraffin embedding. 5um-thick sections were cut and hydrated. Sections were subjected to Masson’s Trichrome staining, according to manufacturer’s instructions or immunohistochemistry by using antibodies listed in table S1. Heat-mediated antigen retrieval was performed before staining. Normal goat serum (10%, Vectorlabs S-1000) was used as blocking buffer. Primary antibodies were visualized with Alexa-Fluor 488, 594, 647 secondary antibodies or with Alexa-Fluor 488- or 647-conjugated streptavidin (Invitrogen, 1:1000) after incubation with biotinylated secondary antibodies (Vectorlabs, 1:500). Streptavidin/Biotin blocking kit (SP-2002, Vectorlabs) was used in case of fluorophore-conjugated streptavidin detection. Nuclei were counter-stained with Hoechst (10mg/ml, diluted to 1:7500). ProLong Gold Antifade (Invitrogen) was used as mounting medium. Negative controls with either no primary antibody or corresponding IgG isotypes.

Bright-field images and Masson’s Trichrome staining were acquired on a Leica DM4000 microscope. Immunofluorescent images were acquired with Leica Thunder Imager, Zeiss LSM700 or LSM900 confocal microscopes. Experimental samples were acquired and processed in the same way. Cell counting, area and signal quantification were performed on multiple ROIs per sample using ZEN (version 2.6 lite) and ImageJ software (version 1.52p). Several photomicrographs of the injury area were obtained at 20x, 40x or 63x magnification using LSM700, LSM900 or Leica Thunder Imager.

Signal quantification was performed on the experimental samples by subtracting the respective background signal from each tissue section and thresholding the fluorescent signal with ImageJ Software (version 1.52p). The area is expressed as percentage of the area of positive signal over the total area of interest. The quantification from the different micrographs within the sections was averaged to give the final value per sample. When quantifying epidermal markers, only the epidermal area was taken into account. Conversely, when quantifying the positive signal in the dermis, only the dermal compartment of the injury area was considered. The quantification of cell number is expressed as percentage of positive nuclei over the total number of nuclei per area of interest. The quantification of the Lef1^+^ units is expressed as average of discrete Lef1^+^ cell clusters across sections of the injury area within the same sample. It included the Lef1^+^ cell clusters in the epidermis and in the hair follicles in the dermis between the amputation plane and the injury tip. The number of the Lef1^+^ placodes indicates the average of Lef1^+^ cell clusters localized in the epidermis only. To assess the pERK antibody and rule out any interspecies variability of antibody binding, the pERK max pixel value/ROI (10^4^ um^2^) was quantified; each dot corresponds to a ROI with its relative pERK max value (0 to 255 pixel values). Although *Acomys* pERK signal is overall higher when compared to *Mus, Mus* samples also possess ROIs with pERK max values similar to the *Acomys* counterparts (i.e. *Mus* at 6hpi), ruling out a potential technical difference in the pERK antibody efficiency across species. The quantification of ERK activation over time between *Mus* and *Acomys* is normalized to the pERK levels at day zero (uninjured) within each species.

### Western Blot

Ear tissue was collected with an 8-mm biopsy punch circumscribing the original 4-mm ear punch at specified time points and lysed in RIPA buffer containing protease inhibitors. Total proteins were quantified using a BCA assay (Thermo Scientific) and equal amounts of proteins were run on a gradient polyacrylamide gel (Mini-PROTEAN pre-cast gels, BioRad). Proteins were transferred to PVDF membranes and membranes were blocked in bovine serum albumin (BSA, 5%) and probed with the following antibodies: pERK (CST # 4370, 1:2000), β-tubulin (Sigma T4026, 1:500). Primary antibodies were detected with HRP-conjugated secondary antibodies and enhanced chemiluminescence (ECL) (Fisher Scientific).

### Transcriptomics

Half-rings of tissue from the proximal punched ear segment were collected from sexually mature and age- and sex-matched *Acomys* for gene expression profiling at 24hpi and D15, after pharmacological ERK inhibition. The tissue was snap frozen, total RNA was extracted in Trizol (Life Technologies) according to manufacturer’s instructions and processed for QuantSeq 3’mRNA-Seq from Lexogen Services (https://www.lexogen.com/services). Briefly, alignment was performed with the Spliced Transcripts Alignment to a Reference (STAR) tool. The RNA-Seq Quality Control package (RSeQC) was run to comprehensively evaluate the RNA-Seq data. RNA integrity and concentration were measured by Bioanalyzer Systems (from Agilent Technologies) and Qubit Fluorometer, respectively. A total of 17 samples were sequenced, distributed as follows: n=4/control 24hpi, n=3/PD 24hpi, n=5/Control D15, n=5/PD D15. Read counts data were processed with iDEP.92 (integrated Differential Expression and Pathway analysis) and differentially expressed genes (DEGs) were identified using DESeq2 (*57*) (FDR cutoff: 0.10).

The Volcano plot shows statistical significance (-Log_10_ adjusted p-value) plotted against log_2_ fold change expression values. Genes downregulated (negative values) and upregulated (positive values) in PD. Color-coded genes were categorized into: cell cycle (green), activated epidermis (orange), matrisome (red).

Gene set enrichment analysis (GSEA) was performed with GSEA software (v4.1.0) and the Molecular Signaling Database (MSigDB, v7.2) (*24, 58*). The upper part of the GSEA plot shows the Enrichment Score (ES) and the normalized ES (NES) (expression dataset vs. ranked list of genes) is indicated together with the False Discovery Rate (FDR) q and nominal p-value. The x axis indicates the genes (vertical black lines) represented in gene sets. The bottom part represents the degree of correlation of genes, positive correlation (red) and negative correlation (blue).

The heatmap of the ERK-dependent matrisome was generated by interrogating the murine ECM atlas (http://matrisomeproject.mit.edu/ecm-atlas/) with the top DEGs (FDR<0.10) of the PD vs. Control dataset at D15. The matrisome is divided into matrisome-associated genes and core matrisome genes. The matrisome-associated division is subcategorized into: secreted factors (pink), ECM-affiliated proteins (dark orange), ECM regulators (light orange). The core matrisome includes: proteoglycans (cyan), ECM glycoproteins (purple) and collagens (cobalt blue).

The heatmaps relative to the ERK-dependent AP-1 target genes were generated by interrogating the Transcription Factor Target (TFT) collection (MSigDB) and TF binding site database by QIAGEN with the top DEGs between control and PD-treated *Acomys*. The heatmaps indicated genes containing at least one AP-1 binding site (TGANTCA) in their promoter region. The genes encoding for transcription factors were identified through the Gene families analysis of the MSigDB gene sets.

To validate the RNA-Seq data, RNA was reverse transcribed using SuperScript IV VILO Master Mix with ezDNAse Enzyme (ThermoFisher Scientific) and SYBR-green quantitative PCR was performed on a Roche LightCycler Multi-well plate 384 with species-specific primers enlisted in table S2, designed using the NIH Primer-Blast tool (*59*). Spiny mouse transcripts were accessed through the SequenceServer BLAST database (*60*). Gene expression was analyzed using delta-delta-Ct method and normalized against *Tbp* housekeeping gene (encoding for TATA-box binding protein).

### Statistics

The normality of residuals was tested to check the Gaussian data distribution and the F-test was used to assess the equality of variances. Differences across groups were evaluated for statistical significance using two-sided Student’s t-test when comparing two groups normally distributed and with equal variances. Unpaired t-test with Welch’s correction was used when comparing two groups normally distributed with unequal variances. Spearman or Pearson bivariate analyses were used for correlation studies. A two-sided Mann-Whitney test was used when comparing two groups not following a normal distribution. Kruskal-Wallis test with Dunn’s post hoc test was used when comparing more than two groups non-normally distributed. One-way ANOVA with Dunnett’s post hoc test was used to compare more than two groups normally distributed. Two-way RM ANOVA with Sidak’s post hoc test was used to compared more than two groups normally distributed over time or distance. Paired analyses were performed with a two-tailed paired t-test. Potential outliers were defined by Grubbs’ test (alpha=0.05). Graphs show mean ± s.e.m. P-values are shown in the graphs. Statistical significance was assessed as ns (not significant), * P<0.05, ** P<0.01, *** P<0.001, **** P<0.0001. Sample size of each experiment is indicated in the Figure legends. All statistical analyses were performed with GraphPad Prism software (v8.1.1).

## Acknowledgements

We thank each and every member of the Bartscherer, Seifert and Bakkers (Hubrecht)’s groups for helpful discussions.

## Author contributions

AT and KB conceived the study; AT, KB and AWS designed the experiments; AT performed the experiments; AT, KB and AWS analyzed the data and wrote the paper.

## Data and materials availability

The RNA-Seq data will be deposited in the GEO database under a specific accession code before acceptance. All data supporting the findings of this study are present in the paper and the Supplementary Materials. Additional data related to this paper may be requested from the authors.

