## Supplementary Materials for "An ERK-dependent molecular switch antagonizes fibrosis and promotes regeneration in spiny mice (*Acomys*)"

### Fig. S1.

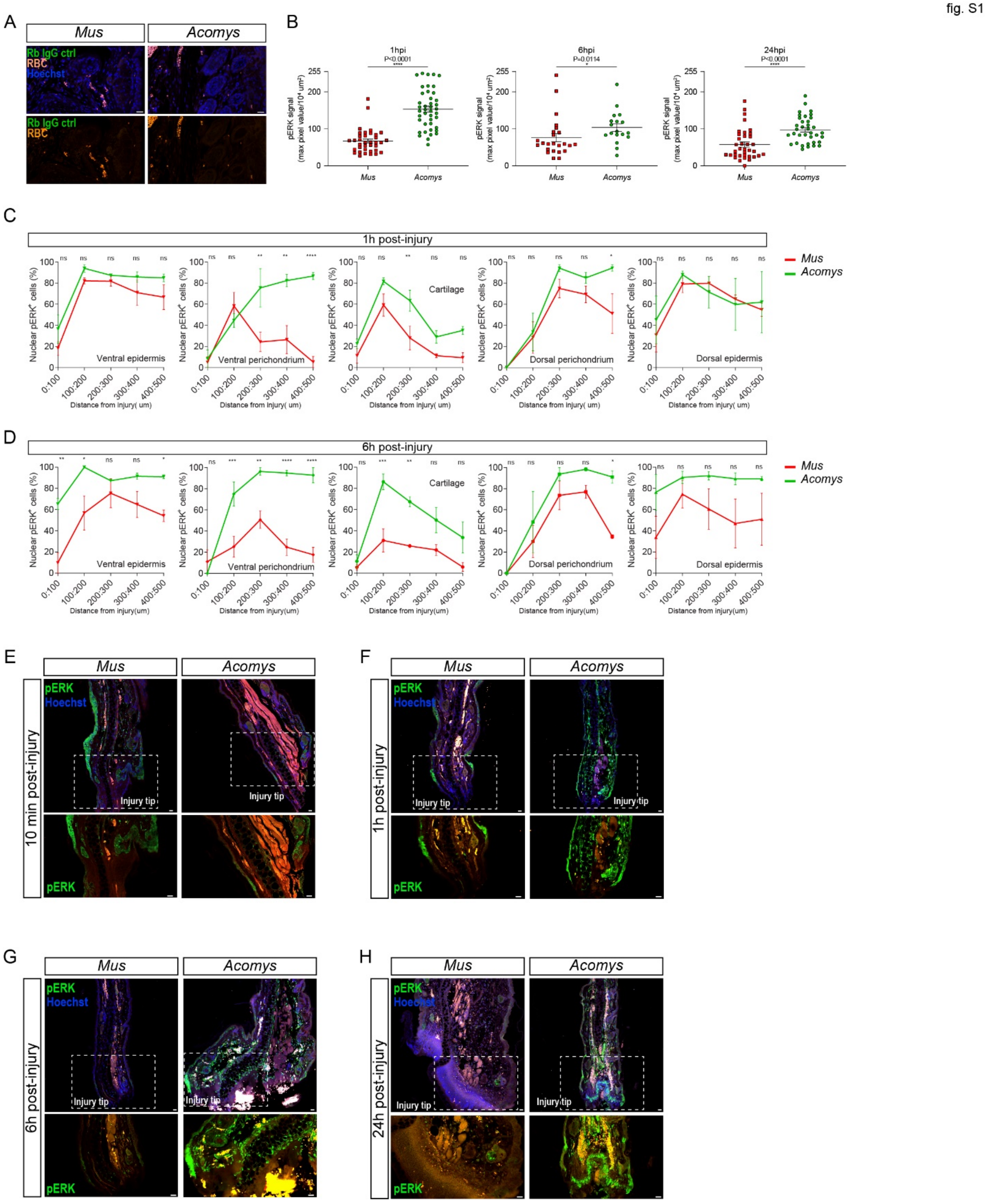

**Fig. S1. Long-distance and long-lasting wave of ERK activity upon injury. (A)** Representative images of IgG negative control in *Mus* and *Acomys* ear sections. **(B)** Quantitation of pERK signal between *Mus* and *Acomys* at 1h post-injury (hpi) (left), 6hpi (central) and 24hpi (right), showing similar antibody reactivity in *Mus* and *Acomys* tissue. Plotted pERK max pixel value/ROI. **(C** and **D)** Quantification of $pERK^+$ nuclei over total nuclei in the area 500um from the injury site in the proximal ear segment at 1hpi (C) and 6hpi (D). **(E to H)**, Representative images of ERK activation (pERK, green) in the distal ear segment between *Mus* and *Acomys* at early time points, zoom-in at the bottom; nuclei (Hoechst, blue). Scale bar: 20um. Two-tailed unpaired t-test with Welch's correction (B, 1hpi), two-tailed Mann-Whitney test (B, 6hpi and 24hpi), Two-way RM ANOVA with Sidak's multiple comparisons test (C and D), n=3-4/species/time point. Data are represented as mean ± s.e.m; ns not significant, * P< 0.05, ** P<0.01, *** P<0.001, **** P<0.0001.

**Fig. S2.**

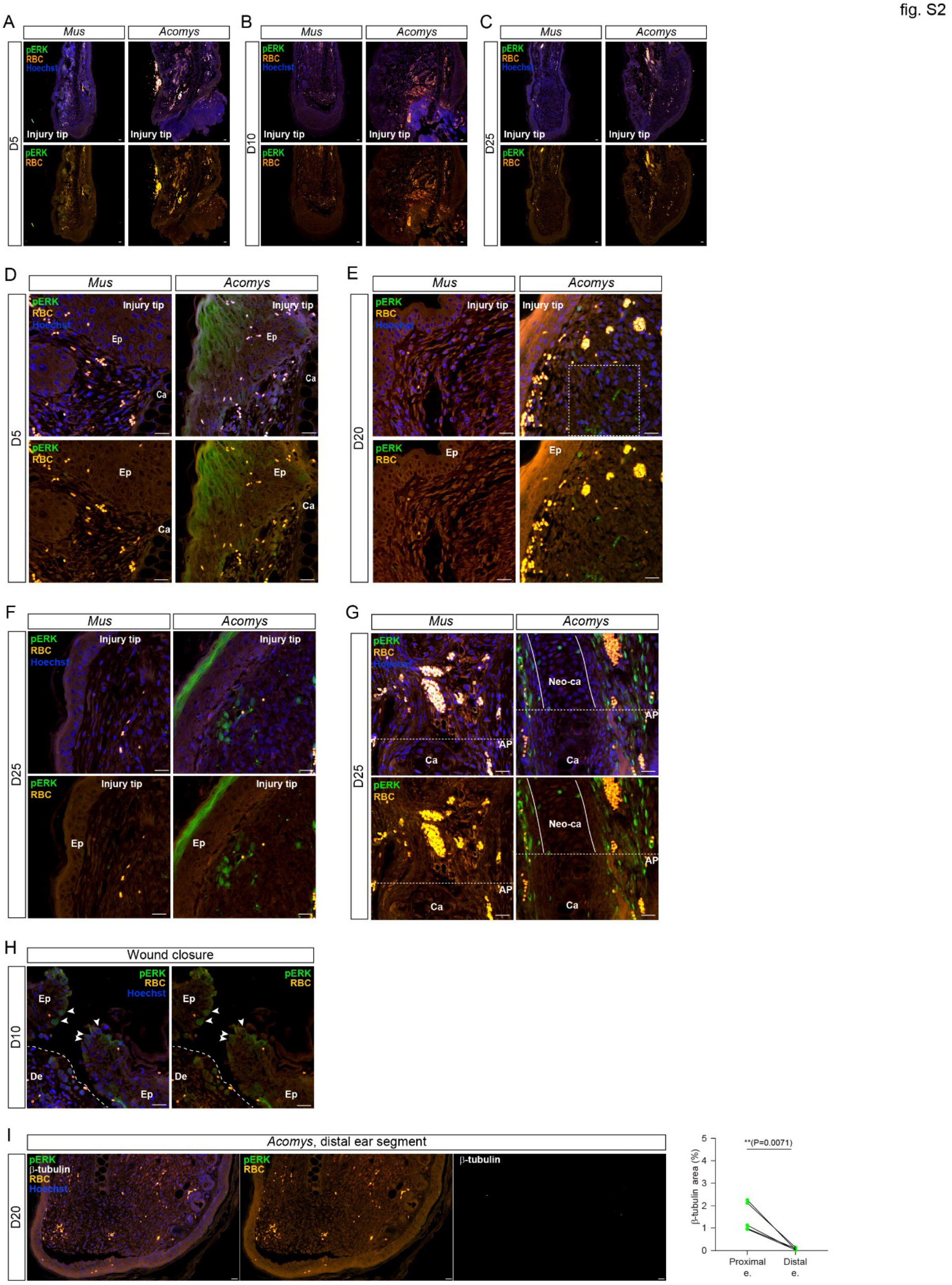
fig. S2
A
B
C
Mus
Acomys
pERK
RBC
Hoechst
Injury tip
D5
D10
D25
D
E
F
G
Ep
Ca
D20
Neo-ca
AP
H
Wound closure
De
I
Acomys, distal ear segment
β-tubulin
β-tubulin area (%)
**(P=0.0071)
Proximal e.
Distal e.
0
1
2
3
4
5

**Fig. S2. ERK activity in the late phase of wound resolution.** (**A** to **C**) Representative images of ERK activation (pERK, green) in the distal ear segment of *Mus* and *Acomys* at late time points, when ERK activation drastically decreases also in distal *Acomys* samples. (**D** to **F**) Representative images of ERK activation (pERK, green) in the proximal ear segment at the injury tip during late time points. **(G)** Representative images of ERK activation at the amputation plane (AP): stream of $pERK^+$ cells along the newly forming cartilage (Neo-ca) in *Acomys*. **(H)** Highlighted $pERK^+$ keratinocytes at the two wound margins undergoing re-epithelialization. Dashed line indicates the epidermis (Ep)-dermis (De) boundaries. **(I)** Representative images of ERK activation (pERK, green) and β-III $tubulin^+$ structures (white) in the distal ear segments (left). Paired analysis of β-III $tubulin^+$ area at the injury tip of proximal and distal ear segments (right). Nuclei counterstained with Hoechst (blue), red blood cells (RBC, orange). Scale bar: 20um. Two-tailed paired t-test (n=5 *Acomys*). Data are represented as mean ± s.e.m;, ** $P<0.01$.

**Fig. S3.**

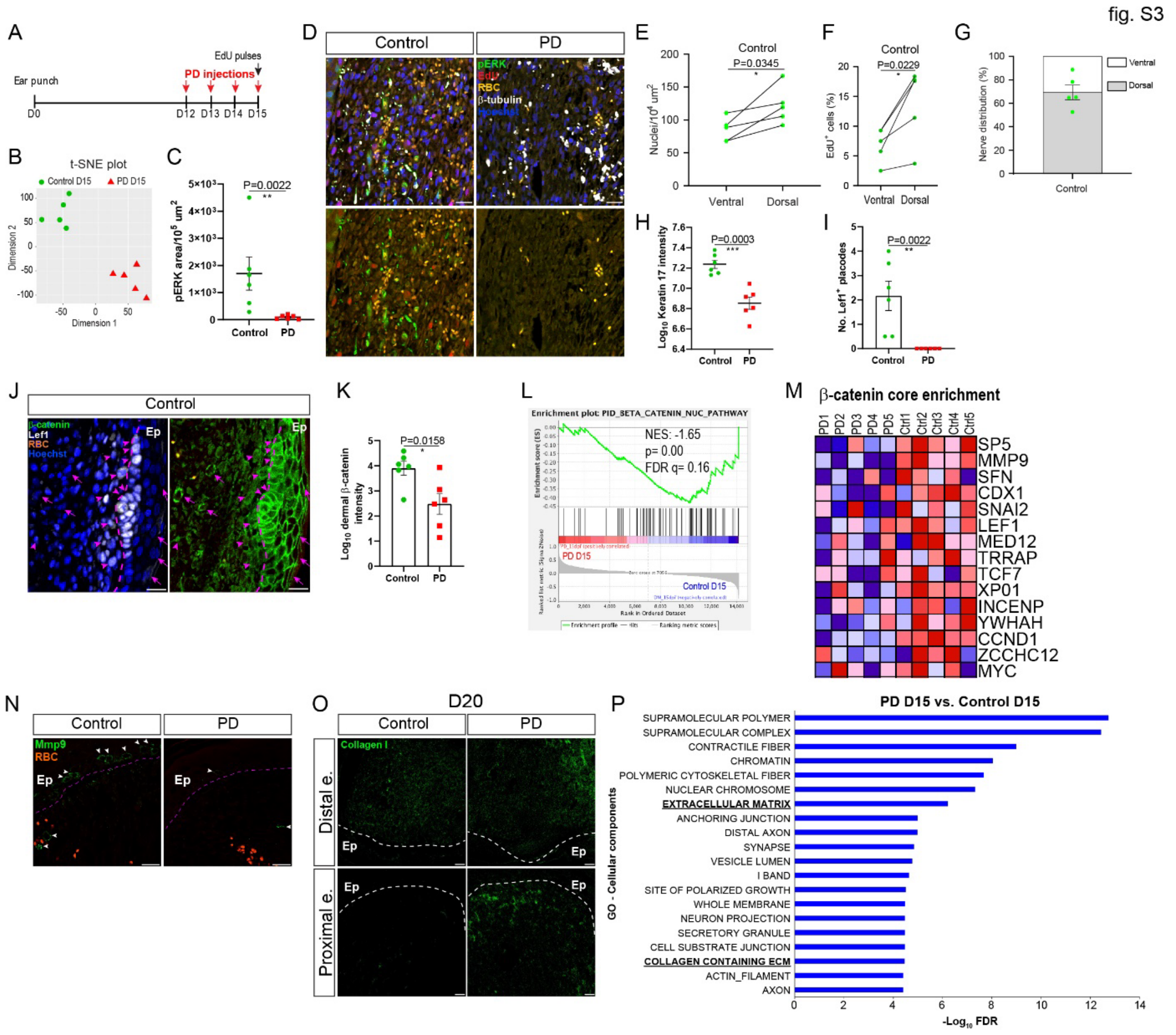


**Fig. S3. ERK inhibition impairs *Acomys* pro-regenerative microenvironment during wound resolution. (A)** Study design. **(B)** t-SNE plot (n=5/group). **(C)** Quantification of ERK activation, D15 (n=6/group). **(D)** Representative images of pERK (green), EdU (red), β-III tubulin (white), D15. **(E** to **G)** Cell density (E), proliferation (F) and nerve distribution (G) across the dorso-ventral axis of the ear pinna. **(H)** Quantification of Keratin17 signal, D15 (n=6/group). **(I)** Quantification of $Lef1^{+}$ hair placodes, D15 (n=6/group). **(J)** Representative images of β-catenin (green) and Lef1 (white), D15 (n=6). Nuclear β-catenin co-localizes with Lef1 (arrowheads); cytoplasmic β-catenin (arrows). **(K)** Quantification of dermal β-catenin signal, D15 (n=6/group). **(L)** GSEA enrichment plot showing the overrepresentation of nuclear β-catenin pathway in the control group and its downregulation in PD group, D15; Normalized Enrichment Score (NES), False Discovery Rate (FDR), nominal p-value (p) (n=5/group). **(M)** Normogram for core enriched genes associated with the β-catenin nuclear pathway. Red and blue represent up- and down-regulation, respectively. **(N)** Representative images of Mmp9, D15. **(O)** Representative images of collagen 1a1 deposition at the injury tip of proximal and distal ear segment. Dashed line, epidermis (Ep)-dermis boundaries. **(P)** Cellular component Gene Ontology analysis. X axis indicates -$Log_{10}$ False Discovery Rate (FDR) statistical significance (n=5/group). RBC (orange), nuclei (Hoechst, blue) Scale bar: 20um. Two-tailed Mann-Whitney test (C and I), two-tailed paired t-test (E and F), two-tailed unpaired t-test (H and K). Data are represented as mean ± s.e.m; ns not significant, * $P< 0.05$, ** $P<0.01$, *** $P<0.001$.

Fig. S4.

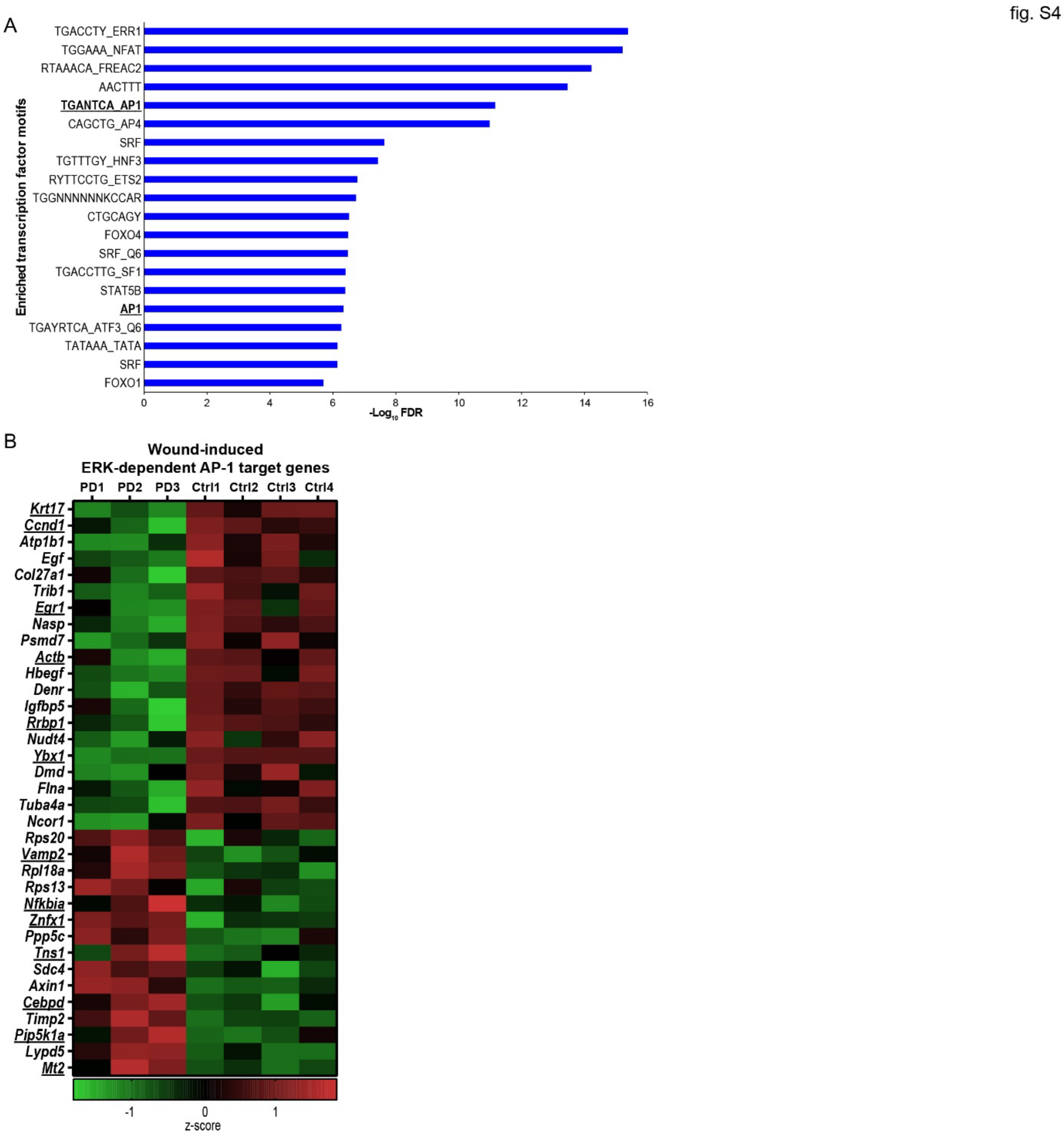


**Fig. S4. Identification of ERK targets during *Acomys* regeneration and early wound response. (A)** GSEA Transcription Factor Targets (TFT) analysis showing the most enriched transcription factor binding sites (y axis) upon ERK inhibition at D15. X axis indicates -Log$_{10}$ False Discovery Rate (FDR) statistical significance (n=5/group). **(B)** Heatmap showing ERK-dependent genes whose promoter contains at least one predicted AP-1 binding site, among the top DEGs between PD and control group expressed at the early wound response (24h post-injury). Underlined are the ERK-dependent DEGs whose expression was induced early by injury and sustained also during the regeneration phase; green and red colors indicate downregulated and upregulated genes, respectively. Gene expression value normalized by z-score transformation

**Fig. S5.**

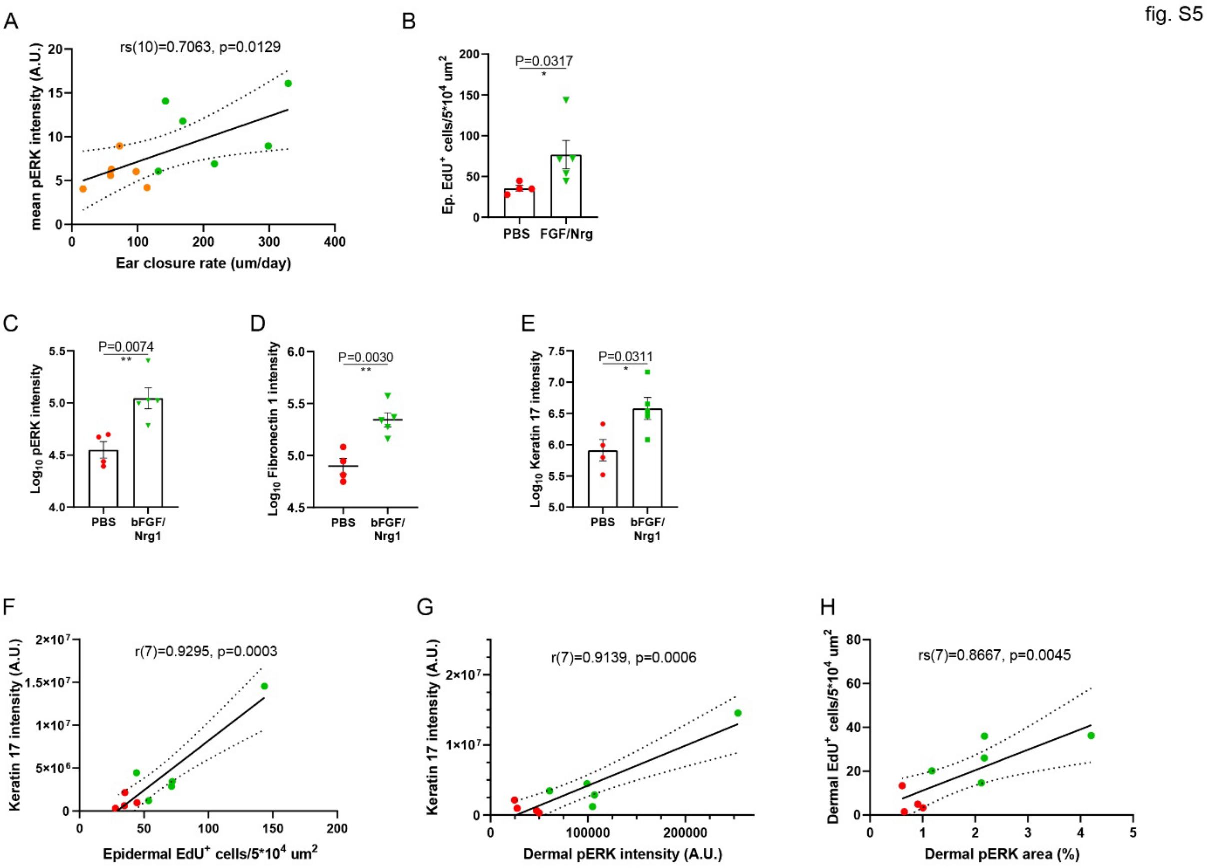


**Fig. S5. ERK activation in hole closure and pro-regenerative effects in normally non-regenerating *Mus* after 4-mm ear punch. (A)** Correlation analysis between ERK activation and closure rate of proximal and distal punches in *Acomys*. Dotted lines indicate 95% confidence intervals; shown Spearman correlation coefficient (rs), degree of freedom (N-2), two-sided p-value. Distal punches (orange), proximal punches (green) (n=6). **(B)** Quantification of proliferating keratinocytes at the injury tip after FGF/Nrg treatment compared to the control group at D20 (n=4 PBS, n=5 FGF/Nrg). **(C** to **E)** Quantification of ERK activation (C), Fibronectin 1 (D), Keratin 17 (E) in punched *Mus* ears treated with FGF/Nrg compared to the control group at D20 (n=4 PBS, n=5 FGF/Nrg). **(F** to **H)** Correlation analysis between Keratin 17 expression and epidermal $EdU^+$ cells (F), Keratin 17 and dermal ERK activation (G), proliferating cells and $pERK^+$ cells in the dermis of punched area (H) in *Mus* at D20. Each dot represents one animal: red dots indicate PBS-treated *Mus*, green dots FGF/Nrg-treated *Mus*. Dotted lines indicate 95% CI (Confidence Intervals); shown Person (r) and Spearman (rs) correlation coefficient, followed by degree of freedom (N-2), two-sided p-value. Two-tailed unpaired t-test (B, C, D and E). Data are represented as mean ± s.e.m; ns not significant, * P< 0.05, ** P<0.01.

**Table S1.**

**Antibody table.**

| Antigen | Host species | Company | Catalog No. |
|---|---|---|---|
| α-smooth muscle actin | mouse | Novusbio | NBP2-34522AF647 |
| β-catenin | mouse | BD Biosciences | 610154 |
| β-III tubulin | mouse | BioLegend | 801201 |
| β-tubulin | mouse | Sigma | T4026 |
| Collagen I | rabbit | Abcam | ab34710 |
| Egr1 | rabbit | Invitrogen | MA5-15008 |
| Fibronectin 1 | rabbit | Sigma | F3648 |
| Keratin 17 | rabbit | Abcam | ab53707 |
| Lef1 | rabbit | Elabscience | E-AB-14187 |
| Mmp9 | rabbit | Abcam | ab38898 |
| PCNA | mouse | Santa Cruz Biotechnology | sc-56 |
| Phospho-ERK | rabbit | CST | 4370 |
| Phospho-histone H3 | rabbit | CST | 9701 |

**Table S2.**

**qPCR primers.**

| Gene | *Mus* ortholog | *Acomys* locus | Forward primer 5'->3' | Reverse primer 5'->3' | Notes |
|---|---|---|---|---|---|
| *Ccl8* | ENSMUSG00000009185 | TRINITY_DN663226 | TGTCATCCCAGGGGAGTTGG | CGCAGATGAGTCTACCACGC | |
| *Ccnb2* | ENSMUSG00000032218 | Locus_451966\|CCNB2_MESAU | CTGCACGCCTCGGGTTTGTA | CCTCCATCCAGAGCAGTTATGGT | |
| *Ccnd1* | ENSMUSG00000070348 | Locus_915836\|CCND1_RAT | CCTCTCCTGCTACCGCACAA | CCTCCTCCTCGGTGACCTTG | |
| *Cd209d* | ENSMUSG00000031495 | Locus_6607\|C209D_MOUSE | ACTGTGCCGAGTTCTCTGGG | ATAGTCTGGATGGCAGGAGTGG | |
| *Fgf12* | ENSMUSG00000022523 | Locus_403123\|FGF12_RAT | AAAGTTCGGGGACACCCACC | GCAGACGAGTTCTCTCGGCTAT | |
| *Mex3a* | ENSMUSG00000074480 | TRINITY_DN1002241 | TGGTGACCGGGAGACGAG | GTTGCGCGAAGCCCTTATC | |
| *Mmp9* | | comp450929_c2_seq2 | TGGTCATGCACTGGGCTTAG | CTTGGGTCAGGCTTAGGGC | ref. (*4*) |
| *Tbp* | | comp453643_c1_seq8 | CTGCGCTGATTTTCAGTTCTGG | AGCTTCTGCACAACCCGA | ref. (*4*) |
| *Tgfbr2* | ENSMUSG00000032440 | Locus_399344\|TGFR2_MOUSE | CCCAAGCCTGACGTGGAACTA | ACTGCACCGCTGTTGTCATTG | |
| *Tgfbr3* | ENSMUSG00000029287 | Locus_392597\|TGBR3_MOUSE | AGTGGACAAGGTGCGGTTCA | TCCAGATCGTGGTGGCATCC | |
| *Vasn* | ENSMUSG00000039646 | Locus_578150\|VASN_MOUSE | GGACGGGCAACTTCTACGAG | ACCTCCCTCTGTTGCCTTGG | |
| *Wisp1* | ENSMUSG00000005124 | Locus_789042\|WISP1_RAT | AGAGGGGCCTGGTTTCACTC | AAACACGGAGCTTGAGCCAC | |
